# Boosting the toolbox for live imaging of translation

**DOI:** 10.1101/2023.02.25.529998

**Authors:** Maelle Bellec, Ruoyu Chen, Jana Dhayni, Antonello Trullo, Damien Avinens, Hussein Karaki, Flavia Mazzarda, Helene Lenden-Hasse, Cyril Favard, Ruth Lehmann, Edouard Bertrand, Mounia Lagha, Jeremy Dufourt

## Abstract

Live imaging of translation based on tag recognition by a single chain antibody is a powerful technique to assess translation regulation in living cells. However, especially in a multicellular organism, this approach is challenging and requires optimization in terms of expression level and detection sensitivity of the system. Here, we improved existing fluorescent tools and developed new ones to image and quantify nascent translation in the living *Drosophila* embryo and in mammalian cells. We tested and characterized five different Green Fluorescent Protein variants fused to the single chain fragment variable (scFv) and uncover photobleaching, aggregation and intensity disparities. Using different strengths of germline and somatic drivers, we determined that the availability of the scFv is critical in order to detect translation throughout development. We introduced a new translation imaging method based on a nanobody/tag system named ALFA-array, allowing the sensitive and simultaneous detection of the translation of several distinct mRNA species. Finally, we developed a largely improved RNA imaging system based on an MCP-tdStaygold fusion.

## Introduction

During development, multipotent cells progressively acquire specific fates leading to the formation of a variety of tissues. This process relies on spatio-temporally regulated gene expression programs, contributing to the reliability and reproducibility of gene expression patterns that generate the final body plan. In this process, the regulation of mRNA translation is a key step since the dynamics of translation in space and time critically affects the outcome of gene expression. Most approaches to study the regulation of translation in whole organisms rely on a bulk population of cells with poor spatial and temporal information. Consequently, the precision and heterogeneity in the regulation of translation at the cellular and molecular level remains incompletely understood. Live imaging methods of translation have been developed in cultured cells^1–5^ since 2016 and recently introduced in *Drosophila*^6–8^. These methods are based on the fluorescent labeling of nascent peptide chains. The labeling system is made of two components, a genetically-encoded antibody such as a single-chain variable fragment antibody (scFv) or Fab fragment^9–11^ fused to a fluorescent protein (FP; referred to as the detector), and a N-terminal peptide epitope inserted in multiple copies in frame with the gene of interest^12^. These live imaging approaches made it possible to visualize translation up to the single-molecule scale in cultured cells but also in an entire organism^7,8^. Such technological breakthroughs provided important insights in our understanding of gene regulation through translation and, for instance, revealed the existence of motor-driven polysome transport and spatial heterogeneity of translation at the subcellular level^1,4,7,13–17^.

The discovery of the *Aequorea victoria* green-fluorescent protein^18,19^ and especially its optimized version mEGFP^20^ have revolutionized imaging of biological processes^21^. Since then, several new FPs have been and continue to be discovered and engineered^22,23^. The constant race for better FPs generally implies the optimization for fast folding, pH resistance as well as improvement of photo-physic properties^24^.

Introduction of the bacteriophage-derived MS2 RNA binding sites, which are recognized by MS2 Coat Protein (MCP) fused to FP, provided a technique to visualize transcription in living cell. This technological advance has been subjected to several improvements from its initial development in 1998^25^. These improvements include increased sensitivity down to single molecules^26^, implementation in a living organism^27^, the generation of new stem loops to visualize single mRNAs at a high temporal resolution^7,28^, artefact-free RNA tags^29^ and double mRNA species imaging with an orthologous system^30^. However, several limitations exist with the present tools that hinder quantitative imaging. For example, protein aggregation has been observed for some systems derived from MS2 Coat Protein fused to GFP^29,31,32^ and for scFv fused to specific GFP variants^12^. Furthermore, the expression level of the detector is critical to ensure accurate detection, particularly for translation, as most of the time one mRNA undergoes multiple rounds of translation. However, no such optimization has been made for the SunTag system in the context of a developing organism, although photo-physical properties of molecules can be variable between different systems (i.e. *in vitro* experiments, cell culture, tissues, organisms…)^24,33,34^.

Here, we tested different GFP variants for their brightness, photobleaching and propensity to aggregation in fixed and live *Drosophila* embryos. We also generated inducible scFv-FP constructs that, when combined with different drivers, extend the use of the SunTag system to a wider range of genes and biological questions. Furthermore, we developed an orthologous method, called ALFA-array, to visualize nascent translation in *Drosophila* embryo and mammalian cells. This system is based on a small and soluble nanobody and we show its usefulness by combining it with the SunTag to visualize translation of two different mRNA species in *Drosophila* embryos.

## Results

### New fluorescent protein fused to scFv to detect nascent translation

The SunTag method is a bipartite system based on an antibody-derived single chain variable fragment (scFv) detector, fused to the GB1 solubility tag and to a fluorescent protein (FP; the detector). The scFv binds a peptide named Supernova Tag (suntag) derived from the yeast GCN4 protein and genetically fused in multiple copies at the N-terminus of the gene of interest^35^ (Figure 1a). To improve the system, we created new scFvs with different green fluorescent proteins under the control of *nanos* (*nos*) enhancer-promoter (EPr). This EPr combination drives expression in the *Drosophila* germline^36–39^ allowing the maternal deposition of mature scFv-FP into laid eggs (Figure S1A). We generated a set of four different constructs, in addition to the previously published scFv fused to the superfolder GFP (sfGFP)^40^ in *Drosophila*, named scFv-sfGFP^7^. The new generation of green fluorescent protein variants includes: i) the monomeric superfolder GFP2 (msGFP2)^41^, ii) the mGreenLantern (GL)^42^, iii) mNeonGreen (NG)^43^ and iv) the monomeric *A. victoria* fluorescent protein 1 (mAvic1)^22^ (Figure 1b, upper panel). Live imaging of these five scFv-FPs during embryonic development until the end of nuclear cycle 14 (n.c. 14) showed that all the scFv-FPs are well expressed, localize mainly in the nucleus due to the presence of a nuclear localization signal, and do not form aggregates (Figure S1b and movie Sup 1-5). Quantification of the fluorescent intensity during the first forty minutes of n.c.14, showed that the apparent brightness of these four new scFv was similar to the scFv-sfGFP, while scFv-mAvic1 produced brighter signals (Figure S1c) and scFv-GL was prone to photobleaching (Figure S1c). We confirmed that the distribution of scFv-FP was homogenous in the embryo and dots/aggregates were not detected by light-sheet imaging of the two brightest scFv-FPs, scFv-mAvic1 and scFv-msGFP2 (Figure S1d and movie Sup 6-7).

**Figure 1:**
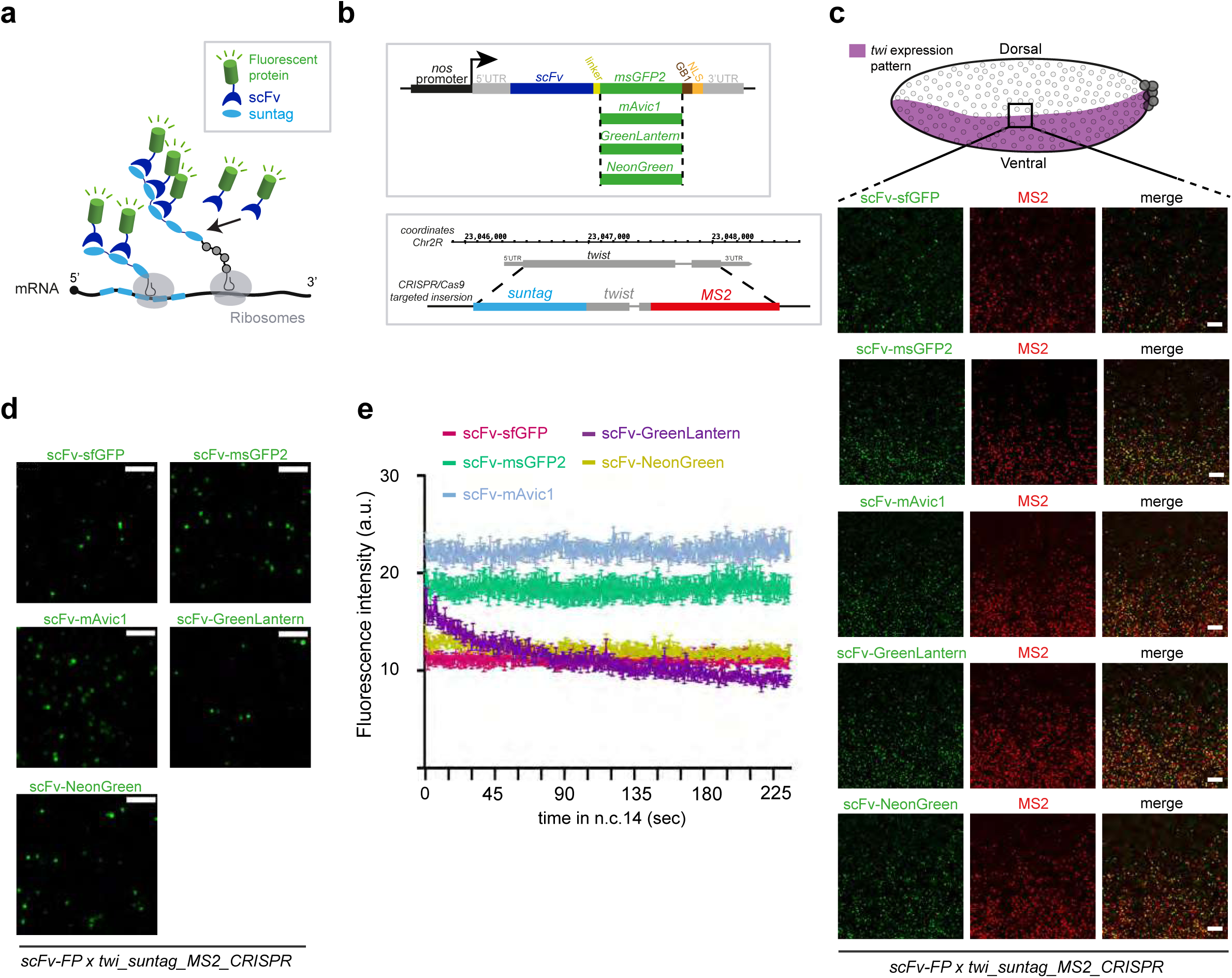
Generation of different scFv-Fluorescent proteins (scFv-FP) to monitor nascent translation in *Drosophila*. **a.** Schematic of the SunTag system. Once the mRNA (black) is translated by the ribosomes (grey), the suntag epitopes (light blue) will be bound by the single chain variable fragment (scFv) (dark blue) fused to a fluorescent protein (FP) (green). Therefore, the nascent peptide will be detected by the accumulation of fluorescent signal. **b.** (top) Schematic of the different constructs of scFv-FP generated in this study. scFv-msGFP2, scFv-GreenLantern, scFv-mAvic1 and scFv-NeonGreen were created for this study and scFv-sfGFP in our previous study^7^. (bottom) Schematic of the CRISPR/Cas9 targeted insertion strategy to obtain endogenous *twist* gene tagged with suntag and 128xMS2^7^. **c.** (top) Schematic of a sagittal view of a *Drosophila* embryo with *twi* expression pattern in purple. Anterior side is on the left, posterior on the right, dorsal on the top and ventral side on the bottom. Black square represents the imaged area of the bottom panels. (bottom) One Z-plane of confocal images from smFISH with direct fluorescent protein signal in green (scFv-msGFP2, scFv-GreenLantern, scFv-mAvic1, scFv-NeonGreen and scFv-sfGFP) and *MS2* probes (red) on *scFv-FP x twi_suntag_MS2_CRISPR* embryos in early-mid n.c. 14. Scale bar are 5µm. **c.** Snapshots from representative fast-mode acquired confocal movies of *twi_suntag_MS2_CRISPR/+* embryos carrying either scFv-msGFP2, scFv-GreenLantern, scFv-mAvic1, scFv-NeonGreen or scFv-sfGFP proteins. Green dots represent nascent translation of *twi*. Scale bars are 5µm. (See related Movies S8-12). **d.** Quantification across time during n.c. 14 of different scFv-FP signals from movies represented in (d), n=5 movies from at least 3 embryos for each scFv-FP. Error bars represent SEM.

We next tested the capacity of these new scFv-FPs to detect mRNA in translation in *Drosophila* embryos. We crossed the different scFv-FP lines into the background of a *twist* gene endogenously tagged with MS2 and suntag (*twi_suntag_MS2_CRISPR*) (Figure 1b, lower panel) and performed single-molecule Fluorescent *in situ* Hybridization (smFISH) on fixed embryos to detect the tagged mRNAs. Actively translated mRNAs (polysomes) were then visualized as co-localized GFP and mRNA spots. Imaging the border of *twist* mRNA transcription pattern showed that the translation signal was present only in the described pattern of *twist* expression (mesoderm)^44^ (Figure 1c), supporting that the signal is specific. For all scFv-FPs, we observed a clear colocalization between mRNA (red signal) and polysomes (green signal) in addition to mRNA molecules that were not detectably translated (Figure S1e). Notably, a fainter signal was detected in fixed samples for polysomes labeled with scFv-mAvic1 at this stage of embryogenesis, as compared to its high apparent brightness in live samples (Figure S1f), suggesting that mAvic1 is more sensitive to the fixation/smFISH protocol. Immuno-smFISH with an antibody against GFP did not improve the signal but increased the background. Surprisingly, we found that staining with only a secondary antibody could label nascent foci of translation (Figure S2a), and only when the scFv-FP was present (Figure S2b). This result suggested that secondary antibodies alone could bind scFv-FP. Experiments using immunofluorescence and scFv-FP simultaneously should thus be avoided. We then looked at the behavior of these scFv-FPs in live embryos in the presence of *twi_suntag_MS2_CRISPR*. We could observe bright green fluorescent dots in movement within the mesoderm (Figure 1d, movie sup 8-12). To characterize the detection capacity of each FP in live embryos, we measured the mean intensity of all detected spots for 4 min during n.c. 14 with a high temporal resolution (Figure 1e, movie sup 8-12). The translation foci detected by scFv-mAvic1, scFv-sfGFP and scFv-msGFP2 showed a stable signal compared to scFv-NG and scFv-GL (Figure 1e). Collectively, these results show that the four new scFv-FPs are able to bind nascent suntag peptides and can therefore detect mRNA being translated in live and fixed samples.

### New scFv-FP prevents aggregate formation

To show whether scFv-FPs expressed under the maternal *nos* EPr could detect translation during late embryogenesis, we imaged fixed embryos at the gastrulation stage. We observed that the tagged Twist protein localized to nuclei in the ventral gastrulation furrow region, consistent with Twist being a mesodermal transcription factor (Figure S2c, upper panel). To test, whether the presence of a NLS in the scFv-FP construct might be responsible for this nuclear localization, we imaged the localization of the Insulin-like peptide 4 (Ilp4), an excreted protein expressed in the mesoderm^45^. We used the tagged Ilp4 with suntag and observed scFv containing the NLS in the cytoplasm (Figure S2c, lower panel). Thus, at least in these cases, the presence of a NLS in the scFv-FP construct appears not to significantly interfere with the native localization of tagged proteins.

Strikingly, at the gastrulation stage, bright GFP spots were observed only when using scFv-sfGFP in *twi_suntag_MS2_CRISPR* embryos but not with the other scFv-FPs (Figure S3a). As only the FP sequences are different between these constructs, this discrepancy might be due to a difference in protein stability, i.e. scFv-sfGFP being more stable than the other variants, or being more prone to aggregation. To distinguish between these possibilities, we used smFISH to label *twi_suntag_MS2_CRISPR* mRNA in the presence of the different scFv-FPs to determine whether the bright GFP spots observed during gastrulation co-localized with mRNA and represent true translation spots. With scFv-sfGFP we saw large green spots that were not co-localized with mRNA (Figure S3b upper left panels, white arrowhead). We did not observe these large spots with any of the four new scFv-FPs (Figure S3b) suggesting an aggregation of the scFv-sfGFP in the *Drosophila* embryo at the gastrulation stage in the presence of *twi_suntag_MS2_CRISPR*.

We next asked whether aggregation was specific for the tagged *twi* in developing embryos or could also occur in cultured cells. We expressed scFv-sfGFP and scFv-msGFP2 in *Drosophila* S2R+ cells along with a SunTag-luciferase reporter (SunTagRLuc-MS2; see methods and Figure S3c). The SunTagRLuc-MS2 plasmid was co-transfected with an scFv-FP plasmid as well as an MCP-Halotag plasmid to track single mRNA molecules and their translation during live-cell imaging. As expected, scFv-FPs were able to label polysomes by forming GFP foci that co-localized with mRNAs (Figure S3d). With the scFv-sfGFP, however, we frequently observed GFP aggregates which were not associated with any mRNA molecules. These aggregates were morphologically distinct from translating foci and were more abundant and bigger than the GFP aggregates observed in gastrulating embryos (Figure S3d-e). This result suggests an aggregation of the SunTag peptides bound by the scFv-sfGFP after being translated and released from polysomes. Although few aggregates were also present in cells expressing scFv-msGFP2 with the SunTagRLuc-MS2 reporter (Figure S3d), this occurred with a significantly lower frequency (∼55% of cells without aggregates compared to ∼15% using scFv-sfGFP) and smaller aggregate size, suggesting that msGFP2 is less prone to aggregate formation.

These results support that the scFv-FPs containing newer generation monomeric FPs are more suitable for translation analysis, as they are less likely to form aggregates compared to sfGFP which is able to form weak dimers^41,46^.

### Monitoring translational arrest during *Drosophila* embryogenesis

The low amount of translation spots observed during gastrulation with scFv-sfGFP suggested that *twi* translation maybe have ceased at this stage of development (Figure S3a-b). However, a translational arrest of *twist* mRNAs has not been described previously. This unexpected observation led us to verify whether *twi* transcripts were undergoing a translational arrest during n.c.14 (Figure 2a), and for this we used the four new scFv-FPs. Interestingly, we found that at mid-late n.c. 14 (Figure 2a), the *twi_suntag_MS2_CRISPR* RNA was mostly translated at the border of the mesoderm (Figure 2b and S4a-c). Using scFv-msGFP2 fluorescence intensity quantification, the percentage of mRNA in translation was slightly lower in the center as compared to the border of the mesoderm (44 ± 4% at the center and 61 ±1% at the border see methods). Strikingly, the polysomes intensities were much lower in the center compared to the mesoderm border, suggesting reduced translation (Figure 2b and S4a-c). A translational arrest was recently described for *Hunchback* (*Hb*) using the SunTag system in the *Drosophila* embryo^8^. In this study the authors showed that after being translated in the entire Hb domain (anterior half of the embryo), *hb* translation became restricted to the border of its expression pattern, similarly to what we observed for *twist*. At least two models can explain this result: i) *twi* translation is actively repressed at first in regions where the RNA and protein expression is highest (as suggested for *hb*^8^), or ii) the amount of scFv-FP becomes limiting during mid-late n.c. 14 at first in regions where the RNA and protein expression is highest and prevents the detection of translation (Figure 2c upper panel). To test these hypotheses, we used the *twi_suntag*_*transgene*, which exhibits a stochastic expression pattern, belated transcriptional activation^7^ and thus lower mRNA level as compared to *twi_suntag_MS2_CRISPR.* If *twi* translation is actively repressed, a similar translational arrest should be observed for the *twi_suntag*_*transgene*. In contrast, if the amount of scFv-FP is limiting, translation of the transgene should be detectable over a longer period of time (Figure 2c). Using live imaging, we observed translation spots of the *twi-suntag*_*transgene* at a stage when *twi_suntag_MS2_CRISPR* translation spots vanished (Figure 2d, S4d and Movie S13-15). The translation dots of the *twi_suntag*_*transgene* colocalized with mRNA dots in late n.c. 14 and gastrulation stage as confirmed with smFISH (Figure S4e). These results suggest that the amount of scFv-FP might be limiting when *twi* translation is monitored from its endogenous locus (*twi_suntag_MS2_CRISPR* allele), preventing detection of translation at gastrulation.

**Figure 2:**
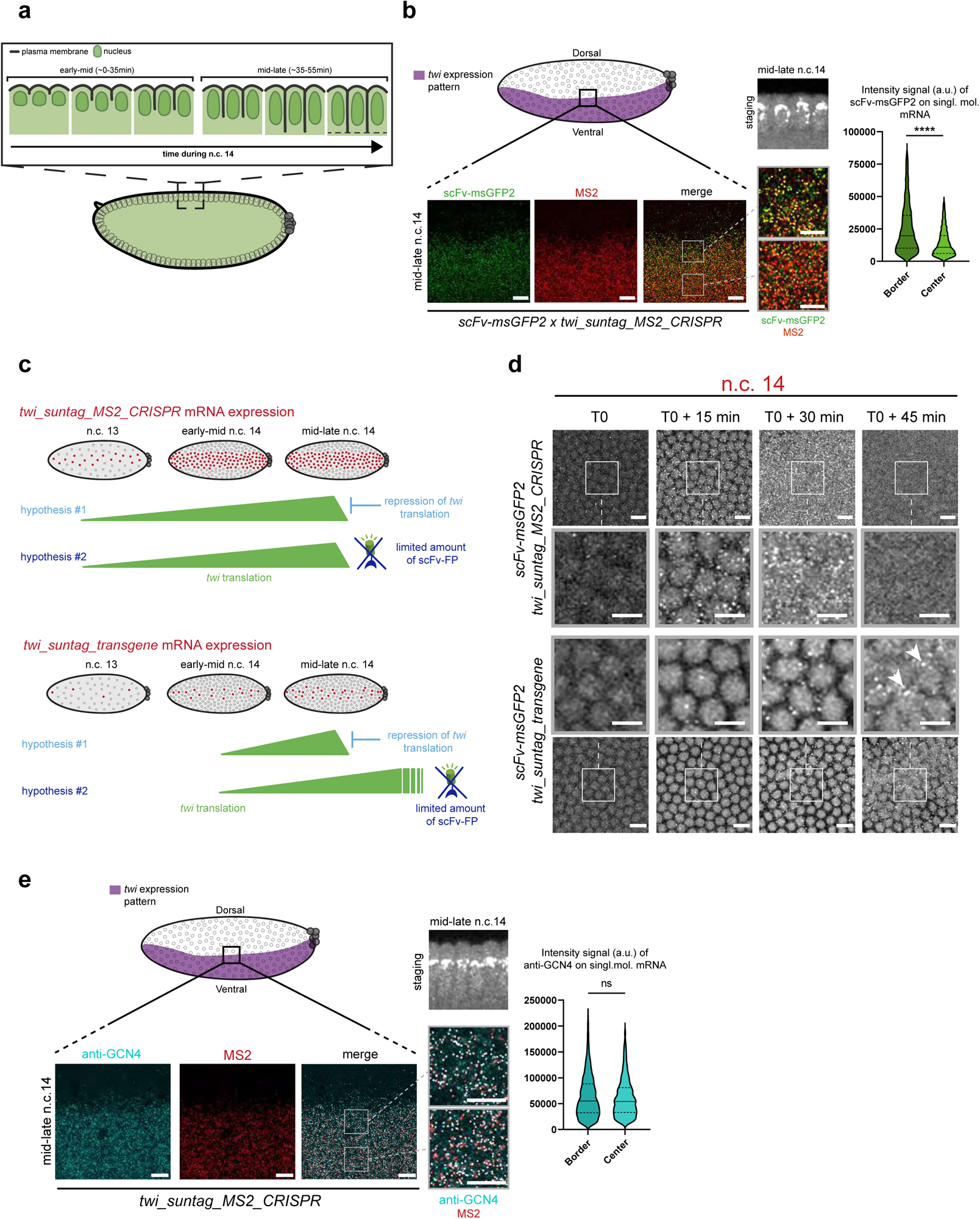
scFv-FP concentration under *nos* EPr creates an artefactual translation arrest of endogenous *twi* mRNA during early embryogenesis. **a.** Schematic of nuclear elongation and cellularization during n.c. 14. In this study, we defined one early-mid stage (corresponding to 0 to 35min of the n.c. 14 from the mitosis) and one mid-late stage (corresponding to 35 to 55min of the n.c. 14 from the mitosis). **b.** (top) Schematic of a sagittal view of a *Drosophila* embryo with *twi* expression pattern in purple. Anterior side is on the left, posterior on the right, dorsal on the top and ventral side on the bottom. Black square represents the imaged area of the bottom panels. (bottom) A single Z-plane of confocal images from smFISH with direct fluorescent protein signal in green (scFv-msGFP2) and *MS2* probes (red) on *scFv-msGFP2 x twi_suntag_MS2_CRISPR* embryos at mid-late n.c. 14. Gray squares represent the zoomed images in the right panels. The two different zoomed images represent border (top) and internal (bottom) zone of the imaged pattern. Scale bars are 10µm on the larger images, and 5µm on the zoomed images. Staging is given by DAPI staining on a sagittal view of the imaged embryo (black and white image at the bottom). Quantification of the scFv-msGFP2 signal intensity on single molecule mRNA at the border at mid-late n.c. 14 stage (dark green, 9 images from three embryos, n=2570) and center (light green, 9 images from three embryos, n=2737) of the mesoderm is represented on the right. **c.** Schematic representing *twi_suntag_MS2_CRISPR* mRNA expression from n.c. 13 to late n.c. 14 in red. From the observations on *twi* CRISPR translation (b), two hypotheses can be considered. Hypothesis 1 is that the decrease of translation observed at mid-late n.c. 14 is due to an active repression of translation. Hypothesis 2 is that the amount of scFv-FP becomes limiting at mid-late n.c. 14, leading to an arrest of the detected green signal. These two hypotheses are represented in the context of *twi_suntag_MS2_CRISPR* expression (above) and *twi_suntag_transgene* expression (below). *twi_suntag_transgene* mRNA is expressed more stochastically and later than *twi_suntag_MS2_CRISPR* mRNA. In hypothesis 1, a decrease of translation should be simultaneously observed for the two constructs (mid-late n.c. 14). In hypothesis 2, no arrest of translation should be observed with *twi_suntag_transgene* as the amount of mRNA is lower. **d.** Snapshots each 15 minutes from movies of *scFv-msGFP2 x twi_suntag_MS2_CRISPR* or *scFv-msGFP2 x twi_suntag_transgene* embryos on the ventral side. T0 correspond to early n.c.14. White squares represent the zoomed images in the center of the panel. Note a persistence of translation for the *twi_suntag_transgene* at T0+45min (white arrowhead), absent in *twi_suntag_MS2_CRISPR* embryos. Scale bars are 10µm on the larger images, and 5µm on the zoomed images. (See related Movies S13-14). **e.** (top) Schematic of a sagittal view of a *Drosophila* embryo with *twi* expression pattern in purple. Anterior side is on the left, posterior on the right, dorsal on the top and ventral side on the bottom. Black square represents the imaged area of the bottom panels. (bottom) Single Z-planes of confocal images from immuno-smFISH with anti-GCN4 antibody (cyan) and MS2 probes (red) on *twi_suntag_MS2* embryos at mid-late n.c. 14. Gray squares represent the zoomed images in the right panels. The two different zoomed images represent border (top square) and internal (bottom square) zone of the imaged pattern. Scale bars are 10µm on the larger images, and 5µm on the zoomed images. Staging is given by DAPI staining on a sagittal view of the imaged embryo (black and white image at the bottom). Quantification of the anti-GCN4 signal intensity on single molecule mRNA at the border (dark blue, 6 images from two embryos, n=3211) and center (light blue, 6 images from two embryos, n=4001) at mid-late n.c. 14 stage of the mesoderm is represented on the right.

To confirm this hypothesis, we used an antibody against the suntag peptides (anti-GCN4) in the absence of scFv-FP and saw translation of *twi_suntag_MS2_CRISPR* mRNAs even at mid-late n.c. 14 and gastrulation stage (Figure 2e and S5a). Moreover, the percentage of mRNA in translation was homogenous within the mesoderm (72 ±4% at the border and 68 ±2% at the center) and polysome intensity between center and border was similar (Figure 2e), in contrast to the results obtained with scFv-msGFP2 (Figure 2b). Altogether these results demonstrate that there is no translational arrest of *twi* mRNAs during n.c.14. Instead, the amount of scFv-FP expressed under the maternally active *nos* EPr becomes insufficient to detect translation during gastrulation, when a large pool of tagged mRNAs is translated. Our experiments suggest that anti-GCN4 antibodies should be used as a control to distinguish between true endogenous repression of translation and mere experimentally-dependent depletion of the labelling reagent.

### Inducible scFv-FP allows imaging of translation during embryogenesis

To overcome the limited amount of scFv-FP and allow the sustained imaging of *twi* translation from egg laying to gastrulation, we used the UAS/Gal4 system to increase the expression of scFv-FP during oogenesis. We generated four lines with the different *scFv-FPs* downstream of a *UASp* promoter (Figure 3a and S6a) and crossed these with the maternal *nos-Gal4* driver (Figure 3b). No aggregation was detected in live embryos (Figure S6a, Movie S16-19 and see methods). To quantify the amount of free scFv-FP using *nosEPr-scFvmsGFP2* or *nos-Gal4>UASp-scFv-msGFP2,* we performed Fluorescence Correlation Spectroscopy (FCS) at early/mid n.c.14 in the mesoderm of embryos. Fluorescence signal corresponding to free scFv-msGFP2 was recorded and fitted with diffusion models (Figure S6b). On average, ∼28 times more free scFv-msGFP2 molecules were available in the confocal volume using *nos-Gal4>UASp-scFv-msGFP2* compared to *nosEPr-scFvmsGFP2* at early/mid n.c.14 (∼710 and ∼25 free scFv-msGFP2 respectively) (Figure S6c). Using this system, we asked if it would allow detection of *twi* translation until late n.c. 14 and gastrulation stage. In *twi_suntag_MS2_CRISPR* embryos with maternally provided *nos-Gal4>UASp-scFv-msGFP2* we saw that *twi* mRNAs were translated throughout n.c.14 (Figure 3c), even in the most ventral part of the embryo, at the center of the *twi* domain of expression (Figure 3c). However, we were not able to detect small dots of scFv-FP, most likely corresponding to individual mature proteins, probably due to the lower signal-to-noise ratio generated by the increased levels of free *scFv-msGFP2*. We also observed translation dots at the gastrulation stage in fixed *twi_suntag_MS2_CRISPR* embryos with *nos-Gal4>UASp-scFv-msGFP2* (Figure 3d left panels) but not with *EPr-scFv-msGFP2* (Figure 3d right panels). Similar results were observed with live imaging, where *twi_suntag_MS2_CRISPR* mRNAs were translated until gastrulation with *nos-Gal4>UASp-scFv-msGFP2* (Figure 3e and Movie S20) but not with *EPr-scFv-msGFP2.* Control embryos lacking *twi_suntag_MS2_CRISPR* mRNAs did not show any translation spot at any stage (Figure 3e and Movie S20). These results indicate that the use of *nos-Gal4>UASp-scFv-msGFP2* would be more suitable to detect translation until gastrulation. The *UASp* promoter drives expression in the germline but is less efficient in somatic tissues^47^. On the contrary, the *UASt* containing *HSP70* promoter drives expression in the somatic tissue^48^. Recently, the *UASt* promoter has been modified to obtain a promoter, called *UASz*, to drive expression in both somatic and germline tissues and being expressed at least four times higher than *UASp*^49^. Due to the high amount of free scFv observed with the *UASp-FP* lines, we generated a new promoter derived from the *UASz*, in which we removed the translational enhancer *IVS-Syn21*^49^ to prevent excessive expression of free scFv-FP and obtain a better signal-to-noise ratio (Figure S7a, b). This new promoter named *UASy* was fused to the *scFv-msGFP2* and crossed with the *nos-Gal4* driver line. As all the other constructs generated in this study, *nos-Gal4>UASy-scFv-msGFP2* alone did not lead to aggregates during early embryogenesis (Figure S7b and Movie S21). Unfortunately, the translation of *twi_suntag_MS2_CRISPR* mRNA at the end of n.c. 14 showed a similar pattern – although less pronounced – as with *scFv-msGFP2* where translation was mostly present at the border of *twi* pattern (Figure S7c), likely due to the weaker expression of the *UASy* compared to *UASp* promoters and subsequent depletion of scFv in the central mesoderm. Maternal expression of the scFv-FP constructs limits our ability to observe translational dynamics at later stages in development. To overcome this, we crossed *UASy-scFv-msGFP2* with an early zygotic *nullo-Gal4* driver to express throughout the embryo. We observed scFv-msGFP2 expression at the beginning of gastrulation (Movie S22) and a strong and homogeneous expression in somatic tissues of later embryos (Figure S7d). We conclude that the *nos-Gal4>UASp-scFv-FP* represents a good tool to image translation from early embryogenesis to gastrulation stage and that the new *UASy-scFv-msGFP2* would enable the imaging of translation in somatic cells at later stages.

**Figure 3:**
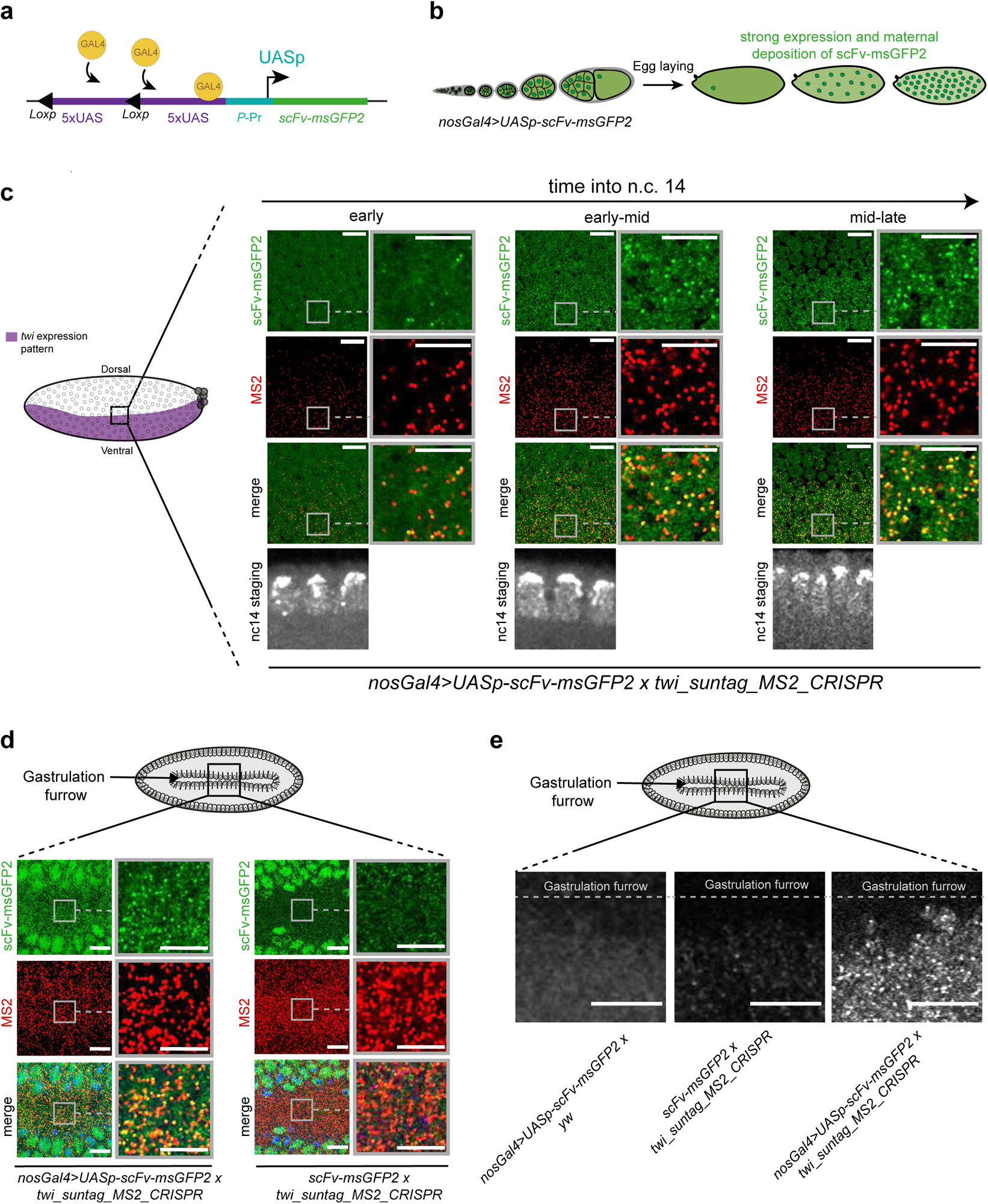
Increasing scFv-FP expression allows translation detection at later stages. **a.** Schematic of *UASp-scFv-msGFP2* construct and activation by the Gal4 protein. 10 UAS sequences (purple) are placed upstream the P-promoter (blue) and scFv-msGFP2 (green) sequences. Gal4 proteins (yellow) will bind the UAS sequences to activate transcription of the scFv-msGFP2. **b.** Schematic of the expression of the *scFv-msGFP2* in *nosGal4>UASp-scFv-msGFP2* female strain during oogenesis until egg deposition. **c.** (left) Schematic of a sagittal view of a *Drosophila* embryo with *twi* expression pattern in purple. Anterior side is on the left, posterior on the right, dorsal on the top and ventral side on the bottom. Black square represents the imaged area of the right panels. Single Z-planes of confocal images from smFISH with direct scFv-msGFP2 fluorescent protein signal (green) and *MS2* probes (red) on *nosGal4>UASp-scFv-msGFP2 x twi_suntag_MS2_CRISPR* embryos in early, early-mid and mid-late n.c. 14. Gray squares represent the zoomed images in for each panel. Nuclei are counterstained with DAPI (grey, bottom images) for staging. Scale bars are 10µm on the larger images, and 5µm on the zoomed images. **d.** (top) Schematic of a *Drosophila* embryo on the ventral side with gastrulation furrow represented with invaginating cells. Black square represents the imaged area of the bottom panels. (bottom) Single Z-planes of confocal images from smFISH with direct scFv-msGFP2 fluorescent protein signal (green) and *MS2* probes (red) on *nosGal4>UASp-scFv-msGFP2 x twi_suntag_MS2_CRISPR* and *scFv-msGFP2 x twi_suntag_MS2_CRISPR* embryos at gastrulation stage and on the ventral side. Scale bars are 10µm on the larger images, and 5µm on the zoomed images. Note that translation dots are visible only with *nosGal4>UASp-scFv-msGFP2*. **e.** (top) Schematic of a *Drosophila* embryo on the ventral side with gastrulation furrow represented with invaginating cells. Black square represents the imaged area of the bottom panels. (bottom) Single Z-planes of confocal images from movie of *scFv-msGFP2 x yw, scFv-msGFP2 x twi_suntag_MS2_CRISPR* and *nosGal4>UASp-scFv-msGFP2 x twi_suntag_MS2_CRISPR* embryos at gastrulation stage (furrow represented with grey dashed line). Scale bars are 10µm. (See related Movie S20).

### Development of the ALFA-array system to monitor translation in *Drosophila*

After having optimized the scFv-FP-based detector, we sought to generate an optimized orthologous system to the SunTag to detect the translation of two different mRNA species in *Drosophila*. To this end, we searched for a highly specific nanobody/tag pair and decided to use the recently developed ALFA-Tag^50^. The ALFA-Tag is a small peptide that harbors several advantages: its synthetic sequence is absent from the proteome of model organisms, its sequence is smaller than that of the SunTag sequence (15 amino acid compared to 19, respectively Figure 4a), it is monovalent, hydrophilic and was effective in *Drosophila* for protein manipulation^51^. Finally, the nanobody anti-ALFA detector (NB-ALFA) is smaller than the scFv (Figure 4b), is quite soluble and displays a very strong affinity for the ALFA-Tag (measured at 26 pM)^50^. Taking advantage of these properties, we multimerized the ALFA-Tag sequences to create a 12x and 32x_ALFA-array. Each ALFA-tag sequence was flanked by a proline to reduce potential influence of neighboring secondary structures, and a spacer of 5 amino acid (GSGSG) to minimize steric hindrance of neighboring peptide binding sites (Figure 4a, c). The NB-ALFA was fused to the msGFP2 and to the GB1 solubilization tag (also used for the scFv) to avoid potential aggregation caused by the multimerization of ALFA-tag sequences. Finally, we generated a *Drosophila* line with the *NB-ALFA-msGFP2* under the control of *nos* EPr (*NB-ALFA-msGFP2)*. The system was also tested in mammalian cells (see below).

**Figure 4:**
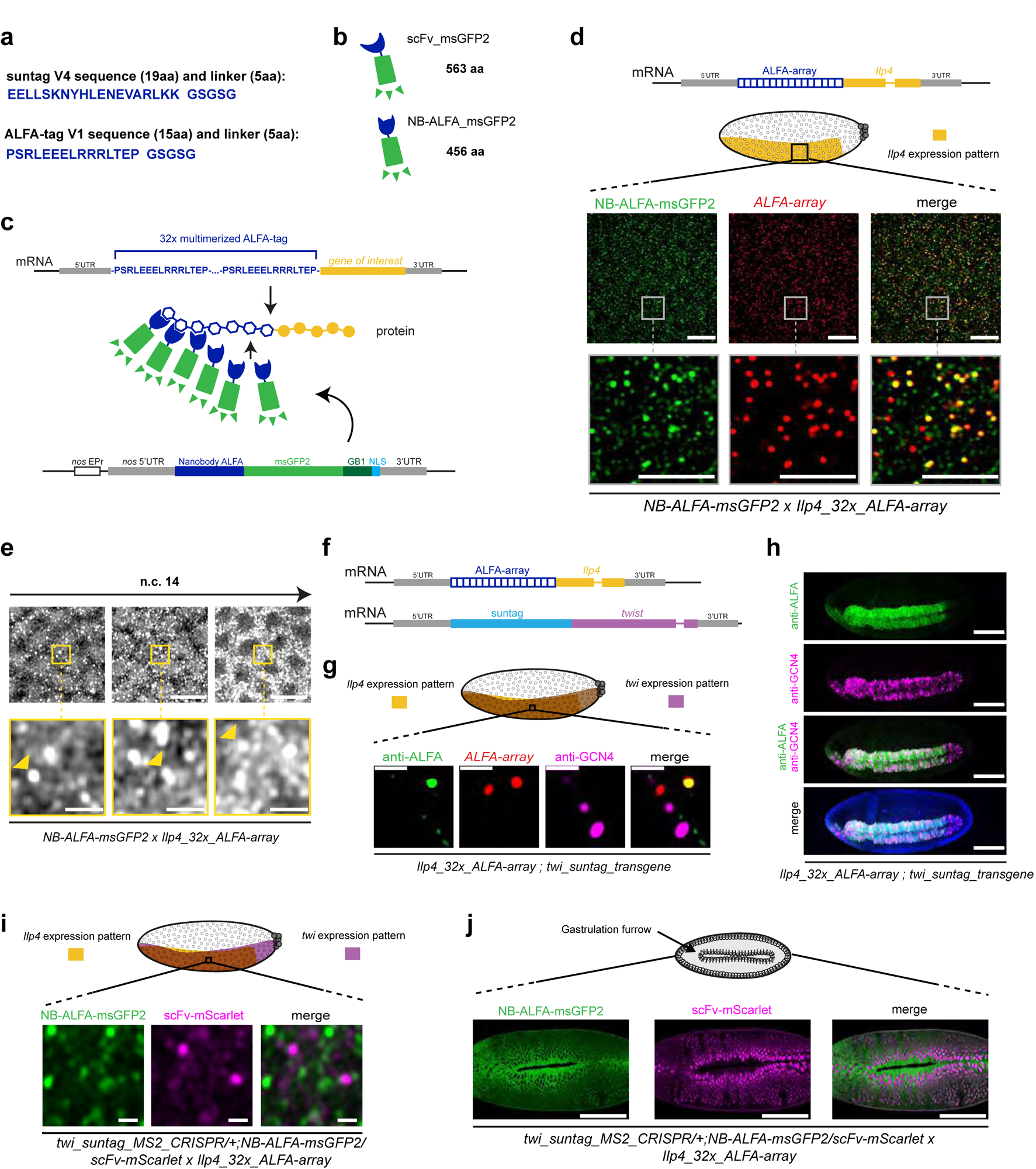
Development of the ALFA-array system to detect nascent translation in *Drosophila*. **a.** Peptide sequence of one suntag with its linker (24 amino-acids) and peptide sequence of the ALFA-tag with its linker (20 amino-acids). **b.** Schematic of the scFv and the nanobody ALFA (NB-ALFA) fused to the fluorescent protein msGFP2 and their size in amino-acid (aa). **c.** Schematic of the ALFA-array system developed in this study to detect translation. 32 ALFA-tag sequences were multimerized (blue amino-acid sequence) and inserted at the 5’ of the gene of interest sequence (yellow). Upon translation, nanobodies ALFA fused to msGFP2 (NB-ALFA_msGFP2) will bind to the ALFA-tag peptides. As a bipartite system, the NB-ALFA is expressed under *nos* EPr fused to *msGFP2*, *Streptococcal protein G* (*GB1*) domain and a *Nuclear Localisation Signal* (*NLS*). **d.** Representation of the transgene mRNA containing 32x_ALFA-array in frame with the *insulin-like peptide 4* (*Ilp4*) gene sequence. Schematic of a sagittal view of a *Drosophila* embryo with *Ilp4* expression pattern in yellow. Anterior side is on the left, posterior on the right, dorsal on the top and ventral side on the bottom. Black square represents the imaged area of the bottom panels. One Z-plane of confocal images from smFISH with direct Nb-ALFA-msGFP2 fluorescent protein signal (green) and 32x_*ALFA-array* probes (red) on *NB-ALFA_msGFP2>Ilp4_32x_ALFA-array* embryos in early-mid n.c. 14 on the ventral side. Scale bars are 10µm on the larger images, and 5µm on the zoomed images. **e.** Snapshots from representative fast-mode acquired confocal movies of *Ilp4_32x_ALFA-array/+* n.c. 14 embryos carrying *NB-ALFA_msGFP2*. Bright white foci represent nascent translation of *Ilp4* transgene. Yellow squares represent zoomed images of the top panels. Nascent translation of *Ilp4* transgene is indicated by yellow arrowheads. Scale bars are 10µm on the larger images, and 2µm on the zoomed images. (See related Movies S24). **f.** Representation of the transgene mRNAs containing 32x_ALFA-array fused to *Ilp4* coding sequence or suntag fused to *twist* coding sequence used in (g) and (h). **g.** Schematic of a *Drosophila* embryo on the ventral side with *twi* expression pattern in purple and *Ilp4* in yellow. Black square represents the imaged area of the bottom panels. One Z-plane of confocal images from immuno-smFISH with anti-ALFA (green) labeling nascent translation of *Ilp4_32x_ALFA-array*, probes against *32x*_*ALFA-array* (red) labeling *Ilp4_32x_ALFA-array* mRNA molecules and anti-GCN4 antibody labeling nascent translation of *twi_suntag_transgene* (magenta) in n.c. 14 embryos. Scale bars are 1µm. **h.** Maximum intensity projection of a whole embryo confocal images from immuno-staining with anti-ALFA antibody (green) and anti-GCN4 antibody (magenta) on *Ilp4_32x_ALFA-array; twi_suntag_transgene* gastrulating embryo. Scale bars are 100µm. **i.** Schematic of a *Drosophila* embryo on the ventral side with *twi* expression pattern in purple and *Ilp4* in yellow. Black square represents the imaged area of the bottom panels. Single Z-plane of confocal images from dual-color live imaging of *twi_suntag_MS2_CRISPR/+; NB-ALFA_msGFP2/scFv-mScarlet x Ilp4_32x_ALFA-array* embryos in n.c. 14. Scale bars are 1µm. (see related Movie S26). **j.** (top) Schematic of a *Drosophila* embryo on the ventral side with gastrulation furrow represented with invaginating cells. Confocal snapshots of a whole live embryo expressing *twi_suntag_MS2_CRISPR/+; NB-ALFA_msGFP2/scFv-mScarlet x Ilp4_32x_ALFA-array*, after gastrulation. Note the cytoplasmic staining from *Ilp4* translation (green) and nuclear staining from *twi* translation (purple). Scale bars are 100µm.

To study translation in *Drosophila* embryos using the 32x_ALFA-array system, we chose to tag the *Ilp4* gene, which codes for a small protein. We reasoned that if we could visualize the translation of a very short transcript/protein, the tool would work even better for longer ones. A 32x_ALFA-array was inserted between the *Ilp4* initiation codon and *Ilp4* gene body, appended with *Ilp4* UTRs and *Ilp4* EPr (*Ilp4_32x_ALFA-array*, Figure 4d). In absence of fluorescent detector expression, we found by smFISH that the *Ilp4_32x_ALFA-array* was expressed in the mesoderm of early *Drosophila* embryos (Figure S8a), consistent with previous reports for *Ilp4*^45^. Moreover, immunofluorescence with the anti-ALFA antibody showed bright dots specifically colocalizing with *Ilp4_32x_ALFA-array* mRNAs and thus representing foci of translation (Figure S8b-c). Importantly, the *Ilp4_32x_ALFA-array* was not recognized by the anti-GCN4 antibody (Figure S8a).

We next confirmed by live imaging that the *NB-ALFA-msGFP2* strain was expressed during embryonic development until the end of n.c.14, that it was mainly localized to the nucleus and that it did not form any aggregates or foci in absence of tagged mRNA (Figure S8d and Movie S23). Then, the *Ilp4_32x_ALFA-array* was crossed with *NB-ALFA*-msGFP2. The resulting embryos showed bright GFP spots overlapping with *Ilp4_32x_ALFA-array* mRNAs (Figure 4d), showing the ability of NB-ALFA-msGFP2 to bind the nascent 32x_ALFA-array *in vivo.* Next, we monitored translation in live embryos and observed bright GFP dots appearing on the ventral side of the embryo (Figure 4e and Movie S24). The signal was restricted to the mesoderm during n.c.14 (Figure S8e and Movie S25) and became more restricted to the ventral furrow during gastrulation (Figure S8f and Movie S25), demonstrating the specificity of the signals. Thus, the *Ilp4_32x_ALFA-array –* NB-ALFA-msGFP2 system is a useful alternative tool to detect mRNA translation during early embryogenesis.

Additionally, staining with a secondary antibody alone could label the translation foci only when the NB-ALFA-msGFP2 is present (Figure S9a-b). This was also observed with the scFv-FPs (see above Figure S2a-b), suggesting that secondary antibodies can directly bind both the NB-ALFA-msGFP2 and the scFv-FP (Figure S2a-b and Figure S9a-b). Knowing that the only common part between NB-ALFA and scFv is the GB1 solubility domain, we tested whether this domain was responsible for binding secondary antibodies. We removed the GB1 from the scFv-msGFP2 and NB-ALFA-msGFP2, expressed them in *Drosophila* S2R+ cells and performed Western-blot analyses (Figure S9c). All the scFv and NB-ALFA constructs (with and without the presence of the GB1) could be detected by an anti-GFP antibody in combination with a suitable secondary antibody. However, when using secondary antibodies alone, the constructs with GB1 were still detected, while no band was observed with the constructs without GB1 (Figure S9c). These results demonstrate the direct binding of the GB1 solubilization domain by secondary antibodies.

To test if we could use the ALFA-array system in combination with the SunTag to image the translation of two distinct mRNA species in the same embryo, the *Ilp4_32x_ALFA-array* was crossed with the *twi_suntag_transgene* (Figure 4f). Using immuno-smFISH in n.c.14 embryos, we observed the two correct expression patterns of Twi and Ilp4, including in the posterior region where only Twi is present (Figure S10a). With higher resolution images, we could visualize single molecules of *Ilp4* mRNA in translation well distinct from *twi* mRNA translation dots (Figure 4g). Furthermore, we were able to distinguish the localization of the Ilp4 protein in the cytoplasm from the Twi protein in the nucleus (Figure 4h). Finally, we then combined four different transgenes, to express scFv-mScarlet, NB-ALFA-msGFP2, *Ilp4_32x_ALFA-array* and *twi_suntag_MS2_CRISPR* in the same embryo. We were able to live image simultaneously the translation of *Ilp4* and *twi* mRNAs (Figure 4i, S10b and Movie S26) in live embryos. Furthermore, we were again able to clearly distinguish the localization of Ilp4 protein in the cytoplasm from Twi protein in the nucleus of live embryos (Figure 4j, S10c and Movie S27). Together, these results demonstrate that the 32x_ALFA-array can be used in conjunction with the SunTag system to study the translation of two different mRNAs during *Drosophila* early embryogenesis in live and fixed samples.

### Development of the ALFA-array system to monitor translation in human cells

To test the performance of the ALFA-tag labeling system to image translation in a different model, we turned to mammalian cells. We first generated HeLa cells expressing a NB-ALFA-sfGFP recombinant nanobody. To evaluate the binding properties of the nanobody for the ALFA tag *in vivo* (measured at 26 pM in vitro^50^) we took advantage of the available H2B_mCherry_ALFAtag constructs^51^. We performed Fluorescence Recovery After Photobleaching (FRAP) of the NB-ALFA-sfGFP, knowing that H2B itself has a very slow recovery time lasting hours^52^. This experiment showed little recovery of NB-ALFA-sfGFP in 5 minutes (Figure S11 a and b), indicating that it is stably bound to its target *in vivo*.

Next, we evaluated the performance of the ALFA-array system to image translation in HeLa cells. We inserted a 12x_ALFA-array repetition at the N-terminus of the DYNC1H1 gene by CRISPR-mediated knock-in (Figure 5a), as previously done for the SunTag^4^. Heterozygous clones were obtained and imaged with and without a thirty-minute puromycin treatment to inhibit translation. In untreated cells, bright foci were observed in the cytoplasm, which disappeared following puromycin addition (Figure 5b). In addition, smiFISH against the endogenous DYNC1H1 mRNAs revealed that the bright ALFA-tag foci colocalized with single mRNAs (Figure 5c), confirming that these corresponded to DYNC1H1 polysomes. The quality of the signals obtained with only 12 repeats of the ALFA-tag demonstrates the efficiency of this labeling system to image translation.

**Figure 5:**
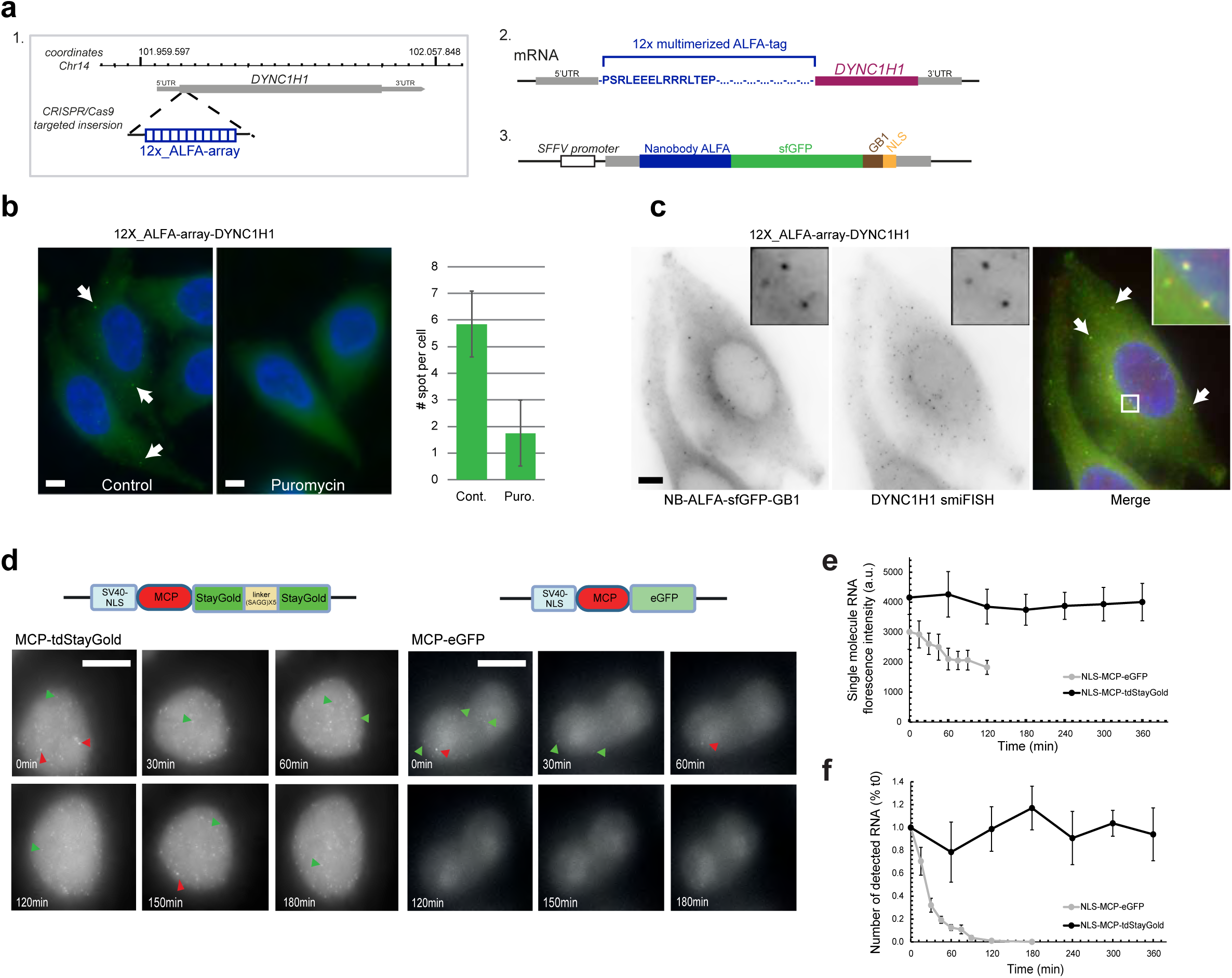
Development of the ALFA-array system to detect nascent translation in mammalian cells. **a.** (1) Schematic of the CRISPR/Cas9 targeted insertion strategy to obtain endogenous *DYNC1H1* gene tagged with 12X ALFA-tag. (2) Schematic of the ALFA-array system developed in mammalian cells. 12 ALFA-tag sequences were multimerized (blue amino-acid sequence) and inserted at the 5’ of the gene of interest sequence (DYNC1H1, yellow). (3) The NB-ALFA is expressed under *spleen focus-forming virus (SFFV)* promoter fused to *sfGFP* and *Streptococcal protein G* (*GB1*) domain. **b.** Left: micrographs of HeLa cells expressing a DYNC1H1 allele endogenously tagged 12x_ALFA-array tag and NB-ALFA-sfGFP, and treated (right) or not (left) with puromycin. Green: NB-ALFA-sfGFP signal; blue: DAPI. Scale bars are 5µm. Arrows: NB-ALFA-sfGFP spots. Right: quantification of the number of NB-ALFA-sfGFP spots per cell, with and without puromycin treatment. Error bars: standard deviation (n=40 cells). **c.** Micrographs of HeLa cells expressing a DYNC1H1 allele endogenously tagged 12x_ALFA-array tag and NB-ALFA-sfGFP, and hybridized in situ with a set of probes against DYNC1H1 mRNAs. On the merged panel: NB-ALFA-sfGFP signal (green); smFISH signals (red); DAPI (blue). Scale bar is 5µm. Arrows: DYNC1H1 polysomes. **d.** Top: schematic of the MCP fusion used to image RNA. Bottom: maximum projection intensity images of cells stably expressing MCP-tdStayGold (on the left) or MCP-eGFP (on the right). All images are taken under the same conditions on an OMX widefield microscope. Six time point are selected (0 min, 30 min, 60 min, 120 min, 150 min and 180 min) and they are displayed with the same dynamic range. Red arrows point at transcription sites and green arrows point at individual RNA molecules. Scale bar: 10 μm. **e.** Intensities of single RNA molecules over time. Graph shows intensities of RNA molecules (mean and s.d. error; five cells). Because of bleaching single molecule signal could no longer be detected and measured after 120 min in the case of MCP-eGFP. Note that MCP-tdStayGold and MCP-eGFP signals were acquired with the same time frame frequency (see methods). **f.** The graph shows the number of single RNA molecule detected over time using FISHquant (five cells).

Finally, an optimal system to visualize RNA is very useful to image translation at the level of single molecules^1–5^. While the MS2/MCP system provides a very nice way to image single RNAs, it is still limited by the low brightness of the signals and its rapid photobleaching. To improve RNA imaging, we took advantage of the recently developed Staygold fluorescent protein, which is bright and exceptionally photostable^53^. Staygold is an obligate dimeric fluorescent protein, and we thus made a tandem dimer version (tdStaygold) by degenerating its sequence and separating two monomers with a (SAGG)x5 linker. This dimeric version of Staygold was then fused to MCP and used to image RNA in a previously developed HeLa cell line expressing an HIV-1 reporter RNAs tagged with 128 MS2 sites^54^. In this system, the MS2 tag is intronic and enables to label nascent RNAs as well nucleoplasmic pre-mRNAs. Live cells were imaged using a frame rate of one 3D image stack every ten seconds for up to 6 hours (Figure 5d). Single RNAs were observed with both NLS-MCP-tdStaygold and the previously used NLS-MCP-eGFP^54^. However, the signal bleached rapidly with NLS-MCP-eGFP, and single RNAs were no longer detectable after 120 minutes. In contrast, the signal showed little photobleaching with NLS-MCP-tdStaygold and single RNAs were efficiently detected through all the movie, and kept their initial brightness (Figure 5e-f). The NLS-MCP-tdStaygold developed here thus shows largely improved performance for RNA imaging.

## Discussion

Amplification of biological signals is crucial to study many biological processes with a high spatio-temporal resolution and single-mRNA-molecule sensitivity. Live imaging of translation at single molecule resolution is a relatively recent technology implemented in 2016 in tissue culture and in 2021 in multicellular organisms. In this study, we provide an expanded toolbox to image translation in living cells and organisms. We will discuss some of the limitations and challenges that need to be considered to obtain robust and reliable results.

Our data show that the choice of the fluorescent protein is a key element for the signal quality of translation imaging-based systems. The development of fluorescent proteins is a very rapidly evolving field and FP-based tools need to be updated to optimize their use. Often, the choice of the FP needs to be empirically defined for each cell type, tissue or organism studied^55–60^. One important aspect in this choice is the monomeric property of the FP to prevent aggregation issues. Although the sfGFP (a weak dimer) was the standard FP for fusion with the scFv in mammalian cell culture, we found that it might present aggregation issue in *Drosophila.* Local translation of specific mRNAs is an important biological process to control gene expression in space and time. To obtain quantitative answers it is necessary to precisely resolve the location of mRNAs and where their translation occurs. In the present experimental designs, with the orthologous scFv and the NB-ALFA, the use of secondary antibodies can confound signal interpretation as they appear to bind the scFv and NB-ALFA fusion. We demonstrate that this binding is due to the GB1 solubilization domain added to limit aggregation and is in line with a recent study in plants, which showed a strong affinity between GB1 and IgG^61^. As the GB1 is required to for the solubility of the detector, combining translation imaging with immunofluorescence must be carried out with great care, and could require a trial with fluorescent primary antibodies.

An important aspect in imaging live embryos or tissues is signal amplification to increase the signal-to-noise ratio. Indeed, in systems like the MS2/MCP, llama-Tag/Fluorescent protein^62^ or scFv/SunTag and their derivatives, the signal appears by local specific accumulation of fluorescent detector, over a diffuse background of molecules freely diffusing in the cells. To improve the signal-to-noise ratio, development of Quenchbody (Q-body) could in theory allow background free detection of translation as well as single mRNA^63^. Currently, the signal-to-noise ratio is optimized by reducing the amount of free fluorescent detector and therefore the background. This is achieved by using weak promoters or targeting the free fluorescent detector to other cellular compartments such as the nucleus, or specific organelles for cytoplasmic analysis. However, one should be careful that the availability of these fluorescent detector does not become limiting as more and more RNAs and proteins are produced during development. As we illustrate here, the available concentration of the detector can be a major challenge in developing organisms, especially when the source of the detector is controlled by an endogenous promoter and when the amount of template mRNA in translation is important. This limitation can also be particularly important when a burst of translation occurs such as during viral infection^14,64^. We provide a direct example for detector limitation during early embryogenesis. Here, early activated genes such as *twist* are highly and rapidly transcribed and translated leading to rapid depletion of free detector. We demonstrate the importance of distinguishing between detector depletion and cessation of translation by validating the results with direct immunostaining. We also show that increasing the expression of the detector can overcome the detection problem, however, it will reduce the signal-to-noise ratio and has thus to be taken into consideration for the specific experimental design. In order to reach conditions that can most accurately capture the dynamics of translation, diverse *UAS* promoters together with a variety of available Gal4 drivers should be tested depending on the tissue of interest and the expression level of the tagged gene.

Studies on mRNA localization and local translation have speculated that different mRNAs species may be translationally co-regulated^65–67^. To obtain independent translational readouts of different mRNA simultaneously, one can combine orthogonal tagging systems such as hybrid scFvs such as HA or FLAG frankenbody^68–70^ or MoonTag nanobodies^16^. In Boersma et al., the authors inserted the MoonTag and the SunTag^16^ into different translational reading frames of the same RNA. This allowed to observe in real time heterogeneities in the start site selection of individual ribosomes on a single mRNA molecule. Here, we introduce a new antibody-based detector/tag pair to image translation in living cells and organisms based on multimerization of the ALFA-Tag. The ALFA-array system has a number of advantages: i) it comes from synthetic design and its sequence is thus absent from organisms; ii) the tag is small, soluble, hydrophilic and lacks aldehydes-sensitive amino acids altered by fixation; iii) the detection is through a nanobody, which is small and soluble, has a very high affinity for the ALFA-Tag (K_d_ of 27 pM), favorably comparable to the MoonTag (Kd of 30 nM)^16^. The new translation imaging system described in this study should be applicable to many different organisms to visualize protein localization and control of translation. Moreover, the availability of several orthogonal systems will allow to better analyze of the complexity of translation dynamics by simultaneously comparing several mRNA in diverse cellular environments and conditions. The wide range of applications of the ALFA-array system, coupled with other antibody-based translation detector, provides to the scientific community a highly versatile toolbox to study translation at the single molecule level in cell lines and multicellular organisms. The combination of nascent protein visualization with improved RNA imaging such as the one provided by the NLS-MCP-tdStaygold will facilitate future scientific breakthroughs.

## Online methods

### Reagents created in this study

| Reagent type | Genotype | Insertion type and chromosome number | Additional information |
| --- | --- | --- | --- |
| *Drosophila* line | *nosEPr > scFv-msGFP2* | Landing site insertion vk33 – Chr. III | mini-white marker, plasmid also available (in pBPhi vector) |
| *Drosophila* line | *nosEPr > scFv-GL* | Landing site insertion vk33 – Chr. III | mini-white marker, plasmid also available (in pBPhi vector) |
| *Drosophila* line | *nosEPr > scFv-mNeonGreen* | Landing site insertion vk33 – Chr. III | mini-white marker, plasmid also available (in pBPhi vector) |
| *Drosophila* line | *nosEPr > scFv-mAvic1* | Landing site insertion vk33 – Chr. III | mini-white marker, plasmid also available (in pBPhi vector) |
| *Drosophila* line | *nosEPr > scFv-msGFP2* | Landing site insertion vk27 – Chr. II | mini-white marker, plasmid also available (in pBPhi vector) |
| *Drosophila* line | *nosEPr > scFv-GL* | Landing site insertion vk27 – Chr. II | mini-white marker, plasmid also available (in pBPhi vector) |
| *Drosophila* line | *nosEPr > scFv-mNeonGreen* | Landing site insertion vk27 – Chr. II | mini-white marker, plasmid also available (in pBPhi vector) |
| *Drosophila* line | *nosEPr > scFv-mAvic1* | Landing site insertion vk27 – Chr. II | mini-white marker, plasmid also available (in pBPhi vector) |
| *Drosophila* line | *UASp > scFv-msGFP2* | Landing site insertion vk33 – Chr. III | mini-white marker, plasmid also available (in pBPhi vector) |
| *Drosophila* line | *UASp > scFv-GL* | Landing site insertion vk33 – Chr. III | mini-white marker, plasmid also available (in pBPhi vector) |
| *Drosophila* line | *UASp > scFv-mNeonGreen* | Landing site insertion vk33 – Chr. III | mini-white marker, plasmid also available (in pBPhi vector) |
| *Drosophila* line | *UASp > scFv-mAvic1* | Landing site insertion vk33 – Chr. III | mini-white marker, plasmid also available (in pBPhi vector) |
| *Drosophila* line | *UASy > scFv-msGFP2* | Landing site insertion - vk33 Chr. III | mini-white marker, plasmid also available (in pBPhi vector) |
| *Drosophila* line | *nosEPr > NB-ALFA-msGFP2* | Landing site insertion - vk33 Chr. III | mini-white marker, plasmid also available (in pBPhi vector) |
| *Drosophila* line | *Ilp4_32x_ALFA-array* | Landing site insertion - vk33 Chr. III | mini-white marker, plasmid also available (in pBPhi vector) |
| DNA plasmid | NB-ALFA in pUC57-Simple | N/A | Synthesized by Genescript, codon optimized |
| DNA plasmid | ALFA-array in pUC57-Simple | N/A | Synthesized by Genescript, with linker GSGSG and codon optimized |
| DNA plasmid | pMT-SuntagRLucFKBP24MS2v5-SV40 | N/A | Used in S2 cell experiments |
| DNA plasmid | pAWF-stdMCP-stdHalotag | N/A | Used in S2 cell experiments |
| DNA plasmid | pAWF-scFv-msGFP2 | N/A | Used in S2 cell experiments |
| DNA plasmid | pAWF-scFv-sfGFP | N/A | Used in S2 cell experiments |
| DNA plasmid | 12x_ALFA-array for CRISPR tagging DYNC1H1 | N/A | Used for HeLa cells experiments |
| DNA plasmid | NB-ALFA-sfGFP in pHR | N/A | Used for HeLa cells experiments |
| DNA plasmid | pAWF-anti-ALFA-msGFP2ΔGB1 | N/A | Used in S2 cell experiments |
| DNA plasmid | pAWF-anti-ALFA-msGFP2 | N/A | Used in S2 cell experiments |
| DNA plasmid | pAWF-scFv-msGFP2ΔGB1 | N/A | Used in S2 cell experiments |
| DNA plasmid | pHAGE Ubc NLS-HA MCP-tdStayGold | N/A | Used for HeLa cells experiments |

### Fly stocks, handling and genetics

The *yw* fly stock was used as a wild type. The germline driver *nos-Gal4:VP16* (BL4937) and the *His2av-mRFP* (Bl23651) fly stocks come from Bloomington Drosophila Stock Center. The *nullo-Gal4* fly stock comes from R.L. laboratory. The *twi_suntag_MS2_CRISPR*, *twi_suntag_transgene* and *scFv-mScarlet* lines were from^7^. Briefly, the 32x suntag sequence from^4^ (containing thirty-three suntag repeats) was inserted in frame into *twi* gene^7^ and 128X MS2 from^28^ placed just after the stop codon. All live experiments were done at 21 °C except when using - or comparing to - strains containing UAS-GAL4 which were done at 25 °C.

### Cloning and transgenesis

*scFv-FP* under *nanos* enhancer-promoter (EPr) lines were generated using NEBuilder® HiFi DNA Assembly Master Mix with primers listed in supplementary table 1 and inserted into scFv-sfGFP vector^7^ digested with BamHI/XhoI. The transgenic construct was inserted into the VK00033 (BL9750) and VK00027 (BL9744) landing site using PhiC31 targeted insertion^71^. *UASp-scFv-FP* lines were done by inserting fragment of the scFv-FP plasmid digested with XhoI/HpaI in the UASp-scFv-sfGFP vector^6^ digested with XhoI/HpaI. The transgenic construct was inserted in the VK00033 (BL9750) landing site using PhiC31 targeted insertion^71^. Fluorescent proteins were extracted from plasmids: mAvicFP1 from pNCST-mAvicFP1 (addgene 129509), mGreenLantern from pcDNA3.1-mGreenLantern (addgene 161912), msGFP2 (addgene 160461), mNeonGreen is a kind gift from S.Piatti. The Ilp4 transgene (regulatory regions and coding sequence) was synthesized (GenScript Biotech) into pUC57-simple^7^. The 32x_ALFA-array repeats were synthesized (GenScript Biotech) into pUC57-simple (see supplementary materials). The 32x_ALFA-array was inserted between Kpn1 and EcoRV restriction sites of the Ilp4 puc57-simple vector. Ilp4-ALFA-array fragment was inserted into pBPhi using PmeI and Nhe1 and injected into BL9750 using PhiC31 targeted insertion^71^ (BestGene, Inc.). The NB-ALFA were synthesized (GenScript Biotech) into pUC57-simple (see supplementary materials), then msGFP2, GB1 and a NLS was added using NEBuilder® HiFi DNA Assembly Master Mix with primers listed in supplementary table 1 and inserted into scFv-sfGFP vector^7^ digested with BamHI/XhoI. The transgenic construct was inserted into the VK00033 (BL9750) landing site using PhiC31 targeted insertion^71^. All cloning for plasmids used in *Drosophila* S2 cells were performed using In-Fusion® HD Cloning Plus Kit (Takara Bio) and PCR was carried out with CloneAmp HiFi PCR premix (Takara Bio). Primers used are listed in supplementary table 1. To inducibly express SunTag in S2 cells, we generated pMT-SuntagRLucFKBP24MS2v5-SV40 by inserting SuntagRLucFKBP24MS2v5 fragment (addgene 119946) into pMT backbone that was PCR-amplified from pMT-EGFPActin5C (addgene 15312) with In-Fusion® HD Cloning Plus Kit. For constitutive expression in S2 cells, pAWF (DGRC stock # 1112) was used as an expression vector. pAWF-stdMCP-stdHalotag was generated by inserting stdMCP-stdHalotag (addgene 104999) into PCR-linearized pAWF vector using In-Fusion® HD Cloning Plus Kit. To generate pAWF-scFv-FPs, sequences containing scFv fused to various fluorescent proteins and GB1-NLS were amplified from *scFv-FP* under *nanos* enhancer promoter plasmids generated in this article and ligated with PCR-linearized pAWF vector using In-Fusion® HD Cloning Plus Kit. All plasmid sequences are listed in supplementary materials. The sequence of anti-ALFA-msGFP2-GB1-NLS was inserted into pAWF vector (DGRC #1112) to generate plasmid pAWF-anti-ALFA-msGFP2. To remove GB1 and generate pAWF-anti-ALFA-msGFP2 ΔGB1, a pair of primers flanking the GB1 sequence were used to amplify the entire pAWF-anti-ALFA-msGFP2 sequence except for the GB1 and ligated with KLD enzyme mix (NEB, M0554S). Same primer pairs were used to generate pAWF-scFv-msGFP2ΔGB1 from pAWF-scFv-msGFP2.

### Single molecule fluorescence *in situ* hybridization (smFISH) and immuno-smFISH in *Drosophila*

A pool of 0-4h after egg-laying (AEL) embryos were dechorionated with bleach for 3 min and thoroughly rinsed with H_2_O. They were fixed in 1:1 heptane:formaldehyde-10% for 25 min on a shaker at 450 rpm; formaldehyde was replaced by methanol and embryos were shaken by hand for 1 min. Embryos that sank to the bottom of the tube were rinsed three times with methanol and kept at -20°C for further use.

Embryos were washed 5 min in 1:1 methanol:ethanol, rinsed twice with ethanol 100%, washed 5 min twice in ethanol 100%, rinsed twice in methanol, washed 5 min once in methanol, rinsed twice in PBT-RNasin (PBS 1x, 0.1% tween, RNasin® Ribonuclease Inhibitors). Next, embryos were washed 4 times for 15 min in PBT-RNasin supplemented with 0.5% ultrapure BSA and then once 20 min in Wash Buffer (10% 20X SCC, 10% Formamide). They were then incubated overnight at 37 °C in Hybridization Buffer (10% Formamide, 10% 20x SSC, 400 µg/ml *E. coli* tRNA (New England Biolabs), 5% dextran sulfate, 1% vanadyl ribonucleoside complex (VRC) and smFISH Stellaris probes against *suntag*^7^ or 32x_ALFA-array coupled to Quasar 570 (supplementary table 1) or probes against 32X MS2 coupled to Cy3 (from E. Bertrand)^54^ and/or GCN4 primary antibody (Novus Biologicals C11L34) (1/200) and/or affinity purified rabbit anti-ALFA polyclonal antibody (NanoTag Biotechnologies) (1/200). Probe sequences are listed in Supplementary table 1. The next day, secondary antibody (1/500 anti-mouse Alexa 488-conjugated (Life Technologies, A21202); anti-mouse Alexa 647-conjugated (Invitrogen, A32728); anti-rabbit Alexa 555-conjugated (Life Technologies, A31572) or anti-rabbit Alexa 488-conjugated (Life Technologies, A21206)) was added if necessary, during 1h at 37° in Wash Buffer. DAPI was added at 45min (1/1000). Embryos were then washed three times 15min in 2x SCC, 0.1% Tween at room temperature before being mounted in ProLong® Diamond antifade reagent.

Images were acquired using a Zeiss LSM880 confocal microscope with an Airyscan detector in Super Resolution (SR) mode. GFP/Alexa-488 was excited using a 488 nm laser, Cy3/Quasar 570 were excited using a 561 nm laser, Alexa 647 was excited using a 633 nm laser. Laser powers were: 27μW for 488nm laser (9μW for UAS>nos images), 80μW for 561nm laser and 46μW for 633nm laser. Laser powers were taken under a 10X objective. Figures were prepared with ImageJ (National Institutes of Health), Photoshop (Adobe Systems), and Illustrator (Adobe Systems).

Percentage of mRNA in translation and polysome intensities were quantified with custom-made algorithms developed in PythonTM^7^. Quantification was done on three area at the center and three area at the border of the mesoderm from three mid-late embryos for *scFv-msGFP2*>*twi_suntag_MS2_CRISPR/+* (related to Figure 2b) and three area at the center and three area at the border of the mesoderm from two mid-late embryos for *twi_suntag_MS2_CRISPR/+* stained with anti-GCN4 antibody (related to Figure 2e). Outliers were removed using the ROUT method^72^, embedded in GraphPad Prism 9.3.1 software, with Q set to 0.1%. scFv-FP intensities (related to Figure S1f) were quantified using images from two to three early-mid *scFv-FP*>*twi_suntag_MS2_CRISPR/+* embryos, outliers were removed using the ROUT method^72^, embedded in GraphPad Prism 9.3.1 software, with Q set to 0.1%.

### Cell culture

Drosophila S2R+ cells (DGRC Stock 150) were maintained at 25°C in Schneider’s medium containing 10% Fetal Bovine Serum and 1% Penicillin-streptomycin. For the transfection, 0.7x10^6 cells in 350ul were seeded into each well of a 24-well plate. Cells are transfected with plasmid mix of pMT-SuntagRLucFKBP24MS2v5-SV40 or pMT-ALFAx32-RLucFKBP24MS2v5-SV40 (200ng), pAWF-MCP-Halotag (5ng), pAWF-scFv-FP-GB1-NLS or pAWF-scFv-FP-NLS (10ng) using effectene transfection reagent. 24h after transfection, cells were first incubated with Schneider’s medium containing 200 nM Janelia Fluor® 594 HaloTag® Ligand (Promega GA1110) for 15min and then switched to medium with 1 mM CuSO4 for 1h or 3h to induce expression of pMT construct.

HeLa cells were maintained in DMEM supplemented with 10% FBS, 10 U/ml penicillin/streptomycin and 2.9 mg/ml glutamine in a humidified CO_2_ incubator at 37°C. For translational inhibition, cells were treated with puromycin at 100 µg/ml for 30 min. Cells were transfected with JetPrime (Polyplus) and selected on 150 µg/ml hygromycin. For each stable cell line, several individual clones were picked and screened by smFISH with sets of fluorescent oligonucleotide probes against the integrated sequence. For CRISPR recombination, clones were additionally analyzed by genomic PCR using amplicons specific for the non-recombined or the recombined allele. The CRISPR guide and repair plasmids have been previously^4^. For the repair plasmid, the 12x_ALFA-array replaced the SunTag 56x tag.

The NB-ALFA-sfGFP was introduced into HeLa cells by retroviral infection. HEK293T cells were transiently transfected with a cocktail of plasmids coding for retroviral components and producing the genomic NB-ALFA-sfGFP retroviral RNAs. Viral particles were collected and used to infect recipient HeLa cells, which were then sorted by FACS. Only lowly expressing cells were selected.

For FRAP experiment, 150k HeLa cells expressing NB-ALFA-sfGFP were seeded in fluorodish 35mm. The day after, cells were transfected with a mix of 3ug of H2B_mCherry_ALFAtag plasmid, 4 µl of Jet Prime Reagent and 200 µl of Jet Prime Buffer. The mix was vortexed for 15 seconds and incubated at RT for 10 minutes before being added to the cells. After 24h, medium containing transfection reagents was washed with PBS and replaced with fresh medium. Cells were incubated for an additional 24h before use. One hour before the FRAP experiment, cells were washed three times with PBS, and incubated in imaging media.

For testing NLS-MCP-tdStaygold, stable expression of fluorescent MCP fusions was achieved by lentivirus-mediated transduction of a self-inactivating vector containing an internal ubiquitin promoter as described in^54^. The MCP contained the deltaFG deletion, the V29I mutation, and an SV40 NLS. Cells expressing fluorescent MCP fusions were grown as a pool of clones and FACS-sorted to select cell populations expressing homogeneous levels of fluorescence. The reporter cell lines additionally expressed the HIV-1 reporter gene pIntro-MS2x128 previously described in^54^.

### Western blot

S2 cells (about 10^6 cells at 3x10^6 cell/ml density) were transfected with 300ng of plasmids pAWF-scFv-msGFP2, pAWF-scFv-msGFP2ΔGB1, pAWF-anti-ALFA-msGFP2, pAWF-anti-ALFA-msGFP2ΔGB1, or empty pAWF vector using Effectene transfection reagent (Qiagen, # 301425). Cells were harvested 48h after transfection, lysed with NuPAGE™ LDS Sample Buffer (Invitrogen, NP0007), and denatured by heating at 70°C for 10 min. Proteins in the lysate samples were resolved by electrophoresis using NuPAGE™ 4 to 12%, Bis-Tris gel, and transferred onto a PVDF membrane using Trans-Blot Turbo Transfer System (Bio-Rad). The PVDF membrane was blocked using Intercept (PBS) Blocking Buffer (LI-COR, 927-70001) for 1 h before being incubated with primary antibodies (rabbit anti-GFP, Invitrogen, A11122; mouse anti-αTubulin, Sigma, T6199-100UL) with 1:3000 dilution overnight at 4°C. When primary antibody was omitted, the membrane was incubated in the blocking buffer overnight at 4°C. The membrane was extensively washed before being incubated with secondary antibodies (Goat anti-Rabbit-HRP, Invitrogen 65-6120; Goat anti-Mouse-HRP, sigma 12-349) at 1:3000 dilution for 1 h at room temperature. The membrane was extensively washed before applying Pierce™ ECL Western Blotting Substrate (Thermo Scientific 32106) and visualized in the BioRad gel imaging system.

### Single molecule fluorescence *in situ* hybridization (smiFISH) in mammalian cells

Cells were grown on glass coverslips (0.17 mm), washed with PBS, fixed in 4% PFA for 20 min, and permeabilized in 70% ethanol overnight at 4°C. Cells were hybridized as previously described for the smiFISH procedure^73^, using probes were sets of unlabeled oligonucleotides hybridizing against the target RNA and additionally contained a common supplementary sequence that was pre-annealed to a fluorescent oligonucleotide probe termed the FLAP^73^. We used sets of oligonucleotides for the DYNC1H1 mRNA^4^. Slides were mounted in Vectashield with DAPI (Vector Laboratories).

Fixed cells were imaged at room temperature on an Axioimager Z1 wide-field microscope (63×, NA 1.4; ZEISS) equipped with an sCMOs Zyla 4 0.2 camera (Andor Technology) and controlled by MetaMorph (Universal Imaging). 3D image stacks were collected with a Z-spacing of 0.3 µm. Figures were prepared with ImageJ (National Institutes of Health), Photoshop (Adobe Systems), and Illustrator (Adobe Systems).

### Live cell imaging of mammalian cells

Cells were plated on FluoroDish (35mm, FD35-100, World Precision Instruments, Sarasota,FL,USA) one day before the imaging. Before imaging media was replaced for imaging purpose with a non-fluorescent media (FluoroBrite DMEM, from thermofischer scientific cat no.A1896701, was supplemented with 10% FBS, 1% penicillin/streptomycin, 1% glutamine and rutin; Evrogen). The dish was mounted in a temperature-controlled chamber with 5 % CO2 and imaged on an inverted OMXv3 Deltavision microscope in time-lapse mode. A 100x, NA 1.4 objective was used, with an intermediate × 2 lens and an Evolve 512 × 512 EMCCD camera (Photometrics). 9 stacks with a z-spacing of 0.6 μm were acquired. Illuminating light and exposure time were set at 1% of laser full power, exposure of 40 ms per plane. For the time-lapse mode, one stack was recorded every 10s for 3h to 6h. Both the MCP-eGFP and MCP-tdStaygold cell lines were imaged under the same conditions.

### Signal quantification in mammalian cells

MCP fusions, quantification was performed using big-FISH, a Python-based software for the analysis of RNA single molecules^74^. Spots were detected using a Laplacian of Gaussian threshold and the program returned the number of the RNA molecule, mean and standard deviation of the fluorescence intensity. The threshold was manually set for each frame for optimal detection of single RNAs and transcription sites were removed from the analysis. Background nucleoplasmic intensities from free fluorescent MCP proteins was subtracted from each frame. The code for the analysis described in this paper is available on GitHub: https://github.com/fish-quant.

### Fluorescence correlation spectroscopy

FCS experiments were performed on a Zeiss LSM780 microscope using a 40x/1.2 water objective. GFP was excited using the 488 nm line of an Argon laser with a constant excitation intensity. Fluorescence intensity fluctuations were acquired for 1 s and auto-correlation functions (ACFs) generated by Zen software were loaded in the PyCorrFit program^75^ and fit using a simple 3D gaussian model:

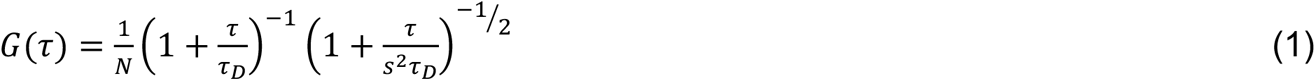

With N the number of molecules, *τ_D_* the diffusion time and s the structural parameter (shape of the PSF) fixed to 5. The brightness B is obtained by dividing the average value of the fluorescence intensity (〈*F*(*t*)〉 by the number of molecules N obtained by the fit of the ACF with equation (1).

Multiple measurements per embryo, in multiple position and embryos at 20 °C were used to generate multiple ACF, used to extract brightness and molecule numbers. Outliers were removed using the ROUT method^72^, embedded in GraphPad Prism 9.3.1 software, with Q set to 0.1%.

### Fluorescence recovery after photobleaching

Fluorescence recovery after photobleaching (FRAP) was performed in HeLa cells expressing NB-ALFA-sfGFP wich was transfected with H2B_mCherry_ALFAtag plasmid^51^ and imaged 48 hours later at 37°C. The following settings was used: a Zeiss LSM980 using a 40x/1.3 Oil objective. Images (512 × 512 pixels), 16bits/pixel, pixel size 0.12µm, zoom 3.5x, were acquired every ≈10 s during 30 frames. Excitation: GFP with a laser at 488nm and mCherry with a laser at 561nm. Laser power was measured using the 40x/1.3 Oil objective at 1mW for 488nm and 561nm laser at 100% and acquisition laser power was measured at 50µW and 2.2µW for 488nm and 561nm diode laser respectively. A circular ROI (3.7µm diameter) was bleached using hundred pulses of 488 nm and 561 lasers at maximal power (100%) after 1 pre-bleach frames. To discard any source of fluorescence intensity recovery other than molecular diffusion, the measured fluorescence recovery in the bleached ROI region (*I_bl_*) was corrected by an unbleached ROI (*I_unbl_*) of the same nucleus and another ROI outside of the cells (*I_out_*) following the simple equation:

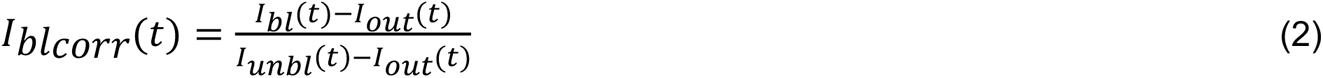

### Light-Sheet Microscopy

For light-sheet imaging (related to Movie S6-7 and Figures S1d), the MuViSPIM^76^ (Luxendo, Brüker company GMBH) was employed. This setup provides two-sided illumination with two Nikon 10x/0.3 water objectives and two-sided detection with two Olympus 20x/1.0 W objectives. The lightsheet is created by scanning a Gaussian beam, as in digital scanned laser light-sheet microscopy (DSLM). We used the line mode which is an in-built synchronization between rolling shutter readout mode of sCMOS cameras and digital light sheets canning. This allows the rejection of out-of-focus and scattered light, thus improving the signal to noise ratio. Images are acquired by two ORCA Flash 4.0 (C91440) from Hamamatsu and processed by LuxControl v1.10.2. A 50ms exposure time was used for the green channel with a 488nm laser. Maximum intensity projections were processed with Fiji^77^.

### Live imaging of the scFv-FP and NB-ALFA-msGFP2 alone

For all live imaging experiments, embryos were dechorionated with tape and mounted between a hydrophobic membrane and a coverslip as described previously^78^. Movies for scFv-FP (related to Figure S1 and Movie S1-5) and NB-ALFA-msGFP2 under *nanos* promoter (related to Figure S8 and Movie S23) were acquired using a Zeiss LSM880 confocal microscope with a Plan-Apochromat 40x/1.3 oil objective lens, with green excitation using a 488nm laser (6μw measured with 10x objective lens). A GaAsP detector was used to detect fluorescence with the following settings: 512x 512-pixels images, zoom 2x, each Z-stack comprised of 25 planes spaced 1μm apart.

Movies for *UASp>scFv-FP* (related to Figure S6, and Movie S16-19) and *UASy>scFv-msGFP2* with *nos-Gal4* driver (related to Figure S7 and Movie S21) were acquired using a Zeiss LSM880 confocal microscope with a Plan-Apochromat 40x/1.3 oil objective lens, with green excitation using a 488nm laser (1μw measured with 10x objective lens). A GaAsP detector was used to detect fluorescence with the following settings: 512x 512-pixels images, zoom 2x, each Z-stack comprised of 25 planes spaced 1μm apart.

### Live imaging of the scFv-FP and anti-ALFA-msGFP2 in presence of tagged genes

For all live imaging experiments, embryos were dechorionated with tape and mounted between a hydrophobic membrane and a coverslip as described previously^78^. Movies of *scFv-FP*>*twi_suntag_MS2_CRISPR/+* (related to Figure 1 and Movie S8-12) were acquired using a Zeiss LSM880 with confocal microscope in fast Airyscan mode with a Plan-Apochromat 40x/1.3 oil objective lens. GFP was excited using a 488nm (7uw measured with 10x objective lens) laser with the following settings: 132x132-pixel images, each Z-stack comprised of 15 planes spaced 0.5μm apart and zoom 8x. Intensity measurement of each polysomes was done on a MIP of 5 planes using a custom-made software written in Python, that performs a spot detection on the MIP raw data stack. For each time frame, the software performs a Laplacian of Gaussian 2D filter and the filtered images are than thresholded with a user defined threshold. The threshold is expressed in terms of the average and standard deviation of intensity values of each image in order to have a threshold value self-re-scaled with respect to the characteristic of the images itself. Spots smaller in size than a user-defined value are removed (considered as fake detection). Finally, for each of the detected spots, we use a 2D expansion of the spots silhouettes in order to define a region around the spots devoided of any other spots and that we can use to estimate background and measure it all along the time evolution for corrections purposes.

Movies of *scFv-FP*>*twi_suntag_MS2_CRISPR/+* and *scFv-msGFP2*>*twi_suntag_transgene* (related to Figure 2, Figure S4 and Movie S13-15) were acquired using a Zeiss LSM880 with confocal microscope in fast Airyscan mode with a Plan-Apochromat 40x/1.3 oil objective lens. GFP was excited using a 488nm (4.7μW measured with 10x objective lens) laser with the following settings: 568x 568-pixel images, each Z-stack comprised of 61 planes spaced 0.5μm apart and zoom 2x. Movies were then processed to remove frame outside of the embryos or containing the membrane signal to correct drifting, and processed stacks were maximum intensity projected using custom-made software, developed in Python^TM7^.

Movies of *nos-Gal4>UASp-scFv-msGFP2* x *twi_suntag_MS2_CRISPR/+* and *scFv-msGFP2* x *twi_suntag_MS2_CRISPR/+* (related to Figure 3e and Movie S20) were acquired using a Zeiss LSM880 with confocal microscope in fast Airyscan mode with a Plan-Apochromat 40x/1.3 oil objective lens. GFP was excited using a 488nm (5uw measure with 10x objective lens) laser with the following settings: 396x 396-pixel images, each Z-stack comprised of 30 planes spaced 0.5μm apart and zoom 4x.

Live tilescans of *nullo-Gal4>*UASy-*scFv-msGFP2-NLS* (related to Movie S22) were acquired using a Zeiss LSM880 with confocal microscope in fast Airyscan mode with a Plan-Apochromat 40x/1.3 oil objective lens. GFP was excited using a 488nm (13μW measured with 10x objective lens) laser with the following settings: 1440 x 2746-pixel images using two tiles, each Z-stack comprised of 9 planes spaced 2μm apart and zoom 0.8x.

Live tilescans of *UASy*-*scFv-msGFP2-NLS/ his2Av-RFP* in presence or not of *nullo-Gal4* driver (related to Supplementary Figure S7d) were acquired using a Zeiss LSM880 with confocal microscope in fast Airyscan mode with a Plan-Apochromat 40x/1.3 oil objective lens. GFP was excited using a 488nm (13μW measured with 10x objective lens) and RFP was excited using a 561nm laser with the following settings: 1024 x 3072-pixel images using three tiles, each Z-stack comprised of 14 planes spaced 2μm apart.

Movies of NB-ALFA-msGFP2 under *nanos* EPr crossed with *Ilp4_32x_ALFA-array* (related to Figure 4e and Movie S24) were acquired using a Zeiss LSM880 with confocal microscope in fast Airyscan mode with a Plan-Apochromat 40x/1.3 oil objective lens. GFP was excited using a 488nm (6.4μW measured with 10x objective lens) laser with the following settings: 256x 256-pixel images, each Z-stack comprised of 6 planes spaced 1μm apart and zoom 6x.

Live tilescans of NB-ALFA-msGFP2 under *nanos* EPr crossed with *Ilp4_32x_ALFA-array* (related to Figure S8e-f and Movie S25) were acquired using a Zeiss LSM880 with confocal microscope in fast Airyscan mode with a Plan-Apochromat 40x/1.3 oil objective lens. GFP was excited using a 488nm (13μW measured with 10x objective lens) laser with the following settings: 1612 x 4860-pixel images using 3 tiles, each Z-stack comprised of 9 planes spaced 2μm apart.

Movies of *NB-ALFA-msGFP2*, *scFv-mScarlet*, *twi_suntag_MS2_CRISPR* crossed with *Ilp4_32x_ALFA-array* (related to Figure 4i and Movie S26) and NB-ALFA-msGFP2, scFv-mScarlet crossed with *Ilp4_32x_ALFA-array* (related to Figure S10b and Movie S27) were acquired using a Zeiss LSM880 with confocal microscope in fast Airyscan mode with a Plan-Apochromat 40x/1.3 oil objective lens. GFP and mScarlet were excited using a 488nm and 561nm laser respectively with the following settings: 372x 372-pixel images, one Z-plane and zoom 6x.

### S2 cell culture imaging

Cells were pipetted into a µ-Slide (Ibidi #80621) and live-imaged using a Zeiss LSM980 confocal microscope with an Airyscan detector in SR mode with a 63X Plan-Apochromat (1.4NA) oil objective lens. GFP was excited using a 488nm laser and MCP-Halotag labelled by Janelia Fluor® 594 was excited using a 561nm laser. Cells with mRNAs visible in cytoplasm and with comparable GFP signal were chosen for acquiring images. Images of individual cells were acquired with following setting: z-stack of 5 planes with 0.5um interval, 8x zoom, 396x396 pixel and 8 bit per pixel. GFP foci co-localized with mRNA foci (Janelia Fluor® 594 signal) were considered as translation sites while GFP foci or larger amorphous or ring-like GFP structures that were not co-localized with mRNA were considered as aggregates.

## Acknowledgments

We are grateful to NanoTag Biotechnologies for sharing ALFA-Tag and anti-ALFA nanobody plasmids and advices. We are grateful to Dr. Karim Mazjoub’s lab for sharing space in his experimental room. We thank Drs. Xavier Pichon and Dr. Vera Slaninova for their help in setting up the mammalian ALFA-Tag system and Dr. Vera Slaninova for helping developing the MCP-tdStaygold fusion. For imaging, we acknowledge Sylvain de Rossi and Montpellier Ressources Imagerie (MRI), a microscopy facility of The National Infrastructure France-BioImaging (FBI), supported by the French National Research Agency (ANR-10-INBS-04). We acknowledge M. Verbrugghe for help with fly handling, and the Montpellier *Drosophila* Facility (BioCampus). We thank Dr. Radhan Ramadass for assistance with software. We acknowledge Dr Etienne Schwob and Dr. Eric Kremer for their help in preparing the manuscript. We acknowledge HFSP (CDA Grant to M.L.) for the early sponsoring of the implementation of *Drosophila* translation imaging project. Funding: This work was supported by the ERC SyncDev starting grant to M.L., the ANR MemoRNP to M.L., and by IGMM internal capital, D.A. was granted by the CNRS consortium GDR ImaBio. M.L., J.D., E.B. and C.F. are sponsored by CNRS. M.B. was supported by a FRM fellowship and then by ERC SyncDev. H.L was supported by ERC SyncDev. J.Dh. was supported by ANR grant ULTIM. E.B. was supported by grants from MSDAvenir and ANR ULTIM.

## Author contributions

J.D. conceived the project. J.D. conceptually supervised the work with the help of M.L. at its early stages. J.D., M.B., M.L., E.B., R.L. and R.C. designed the experiments. J.D., M.B., and C.F. performed *Drosophila* embryos experiments. A.T. developed software. H.L. helped with cloning and fly handling. M.L. acquired funding. E.B. supervised mammalian cells experiments, which were performed by J.Dh.. R.L. supervised *Drosophila* S2 cells experiments performed by R.C.. J.D., M.B., R.C., A.T. E.B. and C.F. analyzed and interpreted the results. C.F. analyzed and supervised FCS experiments with the help of J.D and M.B. J.D. and M.B. wrote the manuscript. M.L., R.C., E.B., R.L. and C.F. edited the manuscript. H.K., F.M. and E.B. developed MCP-tdStaygold fusion. E.B. and J.Dh. developed the mammalian ALFA tag system. J.D., M.B., R.C., D.A., H.K., F.M and J.Dh. performed all experiments done for the review process. All authors discussed and approved the manuscript.

## Competing interests

The authors declare that they have no competing interests.

## Data and materials availability

All relevant data supporting the key findings of this study are available within the article and its Supplementary Information files or from the corresponding author upon reasonable request. The intensity measurement software is available through this link: https://github.com/ant-trullo/Av_Ints_software/. *Drosophila* stocks will be deposited to VDRC stock center and can be asked to M.L. Materials used for S2 cell experiments can be asked to R.L. Materials used for mammalian cell experiments can be asked to E.B.

## Figure legends

**Figure Supp 1:**
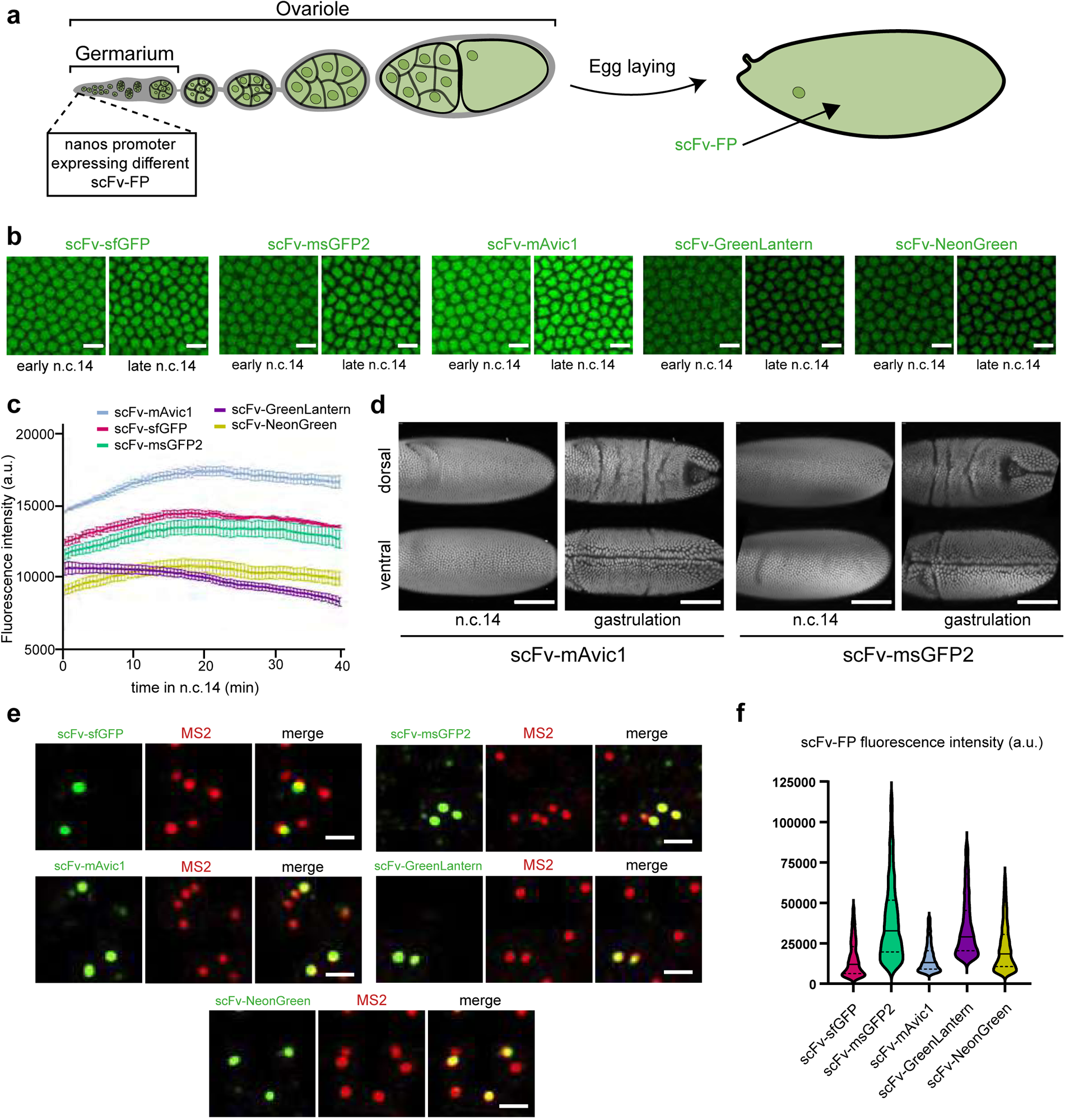
Characterization of the different scFv-Fluorescent Proteins (scFv-FP) **a.** Schematic of the expression pattern of scFv-FP under control of the *nanos* enhancer/promoter during *Drosophila* oogenesis. **b.** Snapshots from movies on embryos carrying scFv-sfGFP, scFv-msGFP2, scFv-GreenLantern, scFv-mAvic1 or scFv-NeonGreen in early and late n.c. 14. Scale bars are 10µm. (see related Movies S1-5). **c.** Quantification across time during n.c. 14 of the different scFv-FP signal from movies represented in (b), n=3 movies from 3 embryos for scFv-sfGFP and scFv-GreenLantern and n=4 movies from 4 embryos for scFv-msGFP2, scFv-mAvic1 and scFv-NeonGreen. Error bars represent SEM. **d.** Snapshots from light-sheet movies on embryos carrying scFv-mAvic1 or scFv-msGFP2 in n.c. 14 and at gastrulation stage. Ventral and dorsal view are represented for each embryo. Scale bars are 100µm. (See related Movies S6-7). **e.** Single Z-planes of confocal images from smFISH with direct fluorescent protein signal in green (scFv-msGFP2, scFv-GreenLantern, scFv-mAvic1, scFv-NeonGreen and scFv-sfGFP) and *MS2* probes (red) on *scFv-FP x twi_suntag_MS2_CRISPR* embryos in early-mid n.c. 14 on the ventral side. Images represent zoomed images from Figure 1c. Scale bars are 1µm. **f.** Quantification of the different scFv-FP signal intensity on single molecule mRNA (n=1646, n=3658, n=3080, n=6820, n=2924, of scFv-sfGFP, scFv-msGFP2, scFv-mAvic1, scFv-GL and scFv-NG respectively) in the mesoderm of early-mid n.c. 14 embryo from immuno-smFISH (cropped area inside *twi* pattern, typical images in Figure 1c).

**Figure Supp 2:**
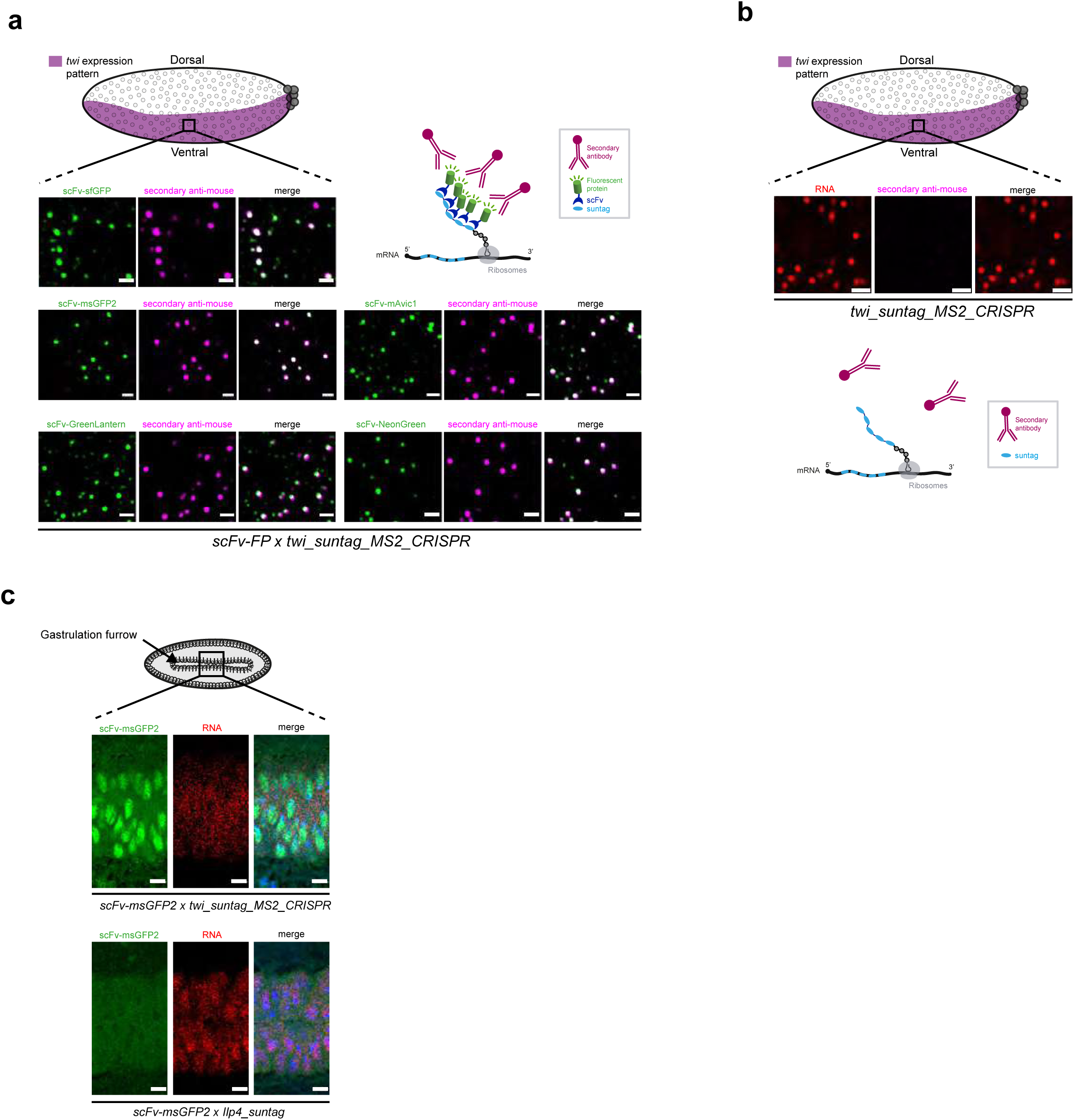
Localization of the tagged protein and scFv recognition by secondary antibody. **a.** (top) Schematic of a sagittal view of a *Drosophila* embryo with *twi* expression pattern in purple. Black square represents the imaged area of the bottom panels. Single Z-planes of confocal images from immuno-smFISH with direct fluorescent protein signal in green (scFv-msGFP2, scFv-GreenLantern, scFv-mAvic1, scFv-NeonGreen and scFv-sfGFP) and secondary anti-mouse antibody (magenta) on *scFv-FP x twi_suntag_MS2_CRISPR* embryos in n.c. 14 on the ventral side. Note the co-localization between the direct fluorescent protein signal and the secondary antibody signal as illustrated by the schematic on the top right. Scale bar are 1µm. **b.** (top) Schematic of a sagittal view of a *Drosophila* embryo with *twi* expression pattern in purple. Black square represents the imaged area of the bottom panels. Single Z-planes of confocal images from immuno-smFISH with MS2 probes (red) and secondary anti-mouse antibody (magenta) on *twi_suntag_MS2_CRISPR* embryos in n.c. 14 on the ventral side. Note that in absence of scFv-FP the secondary antibody does not detect any signal as illustrated by the schematic on the bottom. Scale bars are 1µm. **c.** (top) Schematic of a *Drosophila* embryo on the ventral side with gastrulation furrow represented with invaginating cells. Black square represents the imaged area of the bottom panels. (bottom) Single Z-planes of confocal images from smFISH with direct scFv-msGFP2 fluorescent protein signal (green) and *MS2* or *Ilp4* probes (red) on *scFv-msGFP2 > twi_suntag_MS2_CRISPR* and *scFv-msGFP2 > Ilp4_suntag* embryos at gastrulation stage and on the ventral side. Note a nuclear signal in *twi_suntag_trangene* embryos and a cytoplasmic signal in *Ilp4_suntag* embryos. Scale bars are 10µm.

**Figure Supp 3:**
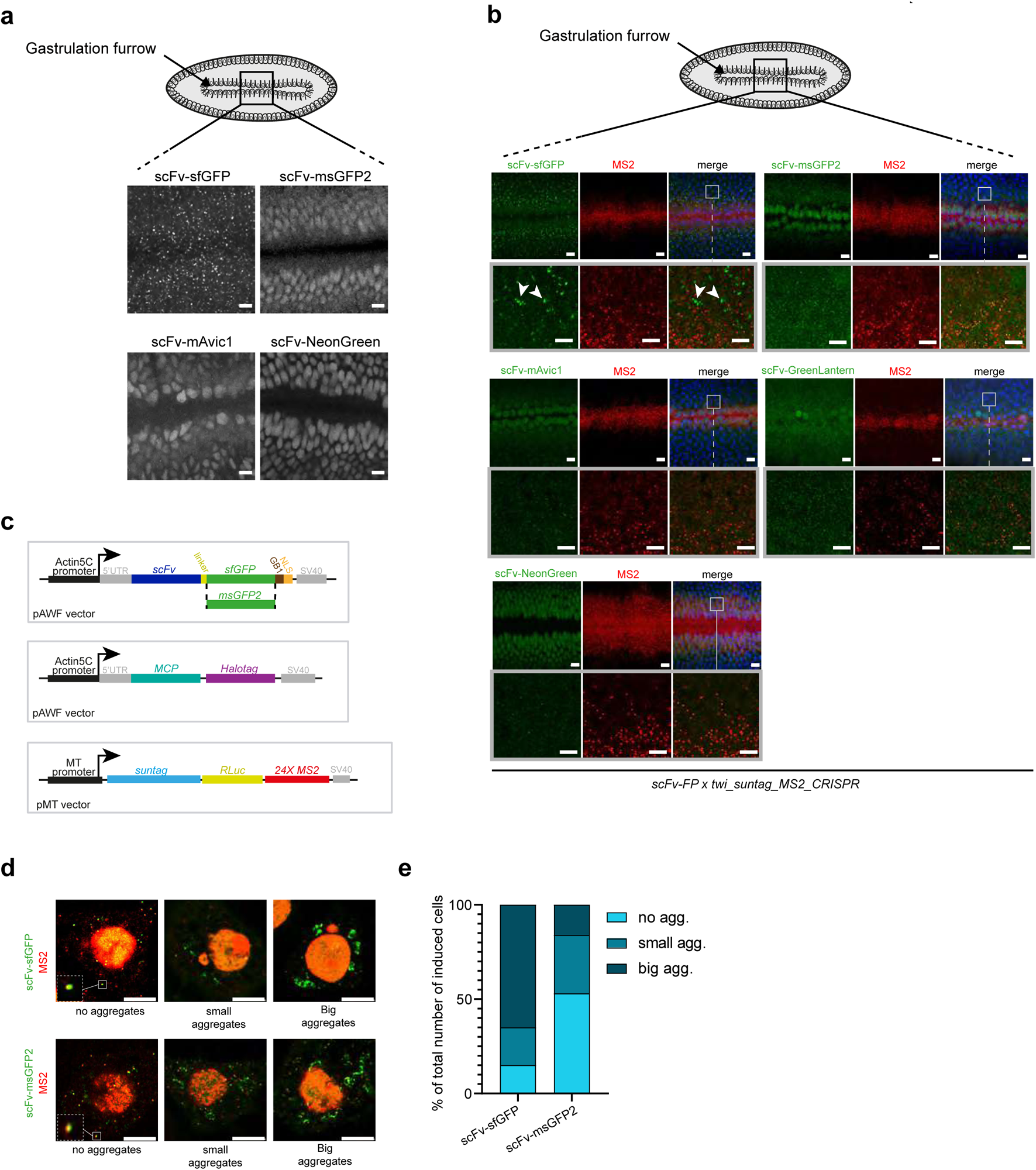
The scFv-sfGFP can form aggregates in *Drosophila* embryos and S2 cells. **a.** Schematic of a *Drosophila* embryo on the ventral side with gastrulation furrow represented with invaginating cells. Black square represents the imaged area of the bottom panels. Snapshots from representative confocal movies of *twi_suntag_MS2_CRISPR/+* embryos at the gastrulation stage carrying either scFv-sfGFP, scFv-NeonGreen, scFv-mAvic1 or scFv-msGFP2 proteins. Scale bars are 10µm. **b.** Schematic of a *Drosophila* embryo on the ventral side with gastrulation furrow represented with invaginating cells. Black square represents the imaged area of the bottom panels. Maximum intensity projection of 10 Z-planes of confocal images from smFISH with direct fluorescent protein signal in green (scFv-FP) and *MS2* probes (red) on *scFv-FP x twi_suntag_MS2_CRISPR* embryos at gastrulation stage on the ventral side. Grey squares represent zoomed images. Note the presence of scFv-sfGFP signal not overlapping with RNA (white arrowheads). Scale bars are 5µm. **c.** Schematic of the different constructs transfected in S2 cells. *scFv-FPs* and *tdMCP-stdHalotag* were placed downstream of the *Actin5C* promoter. Reporter containing suntag Renilla Luciferase (RLuc) and 24XMS2 was placed under the control of the inducible metallothionein (MT) promoter. **d.** Single Z-planes of confocal images of S2 cells expressing scFv-FPs (green) and MCP-Halotag (red) with induction of *SuntagRLuc24XMS2* reporter showing aggregation size of the scFv-FPs. A zoom on single mRNA in translation is shown on no aggregate’s panels (white squares). Scale bars are 5µm. **e.** Percentage of the number of big (dark blue) and small (blue) aggregates (scFv-FP signal not colocalizing with an mRNA) and no (light blue) aggregates (scFv-FP signal colocalizing with an mRNA) in cells expressing *MCP-Halotag*, *SuntagRLuc24XMS2* reporter and *scFv-sfGFP2* or *scFv-msGFP2* (n=100 cells for each condition).

**Figure Supp 4:**
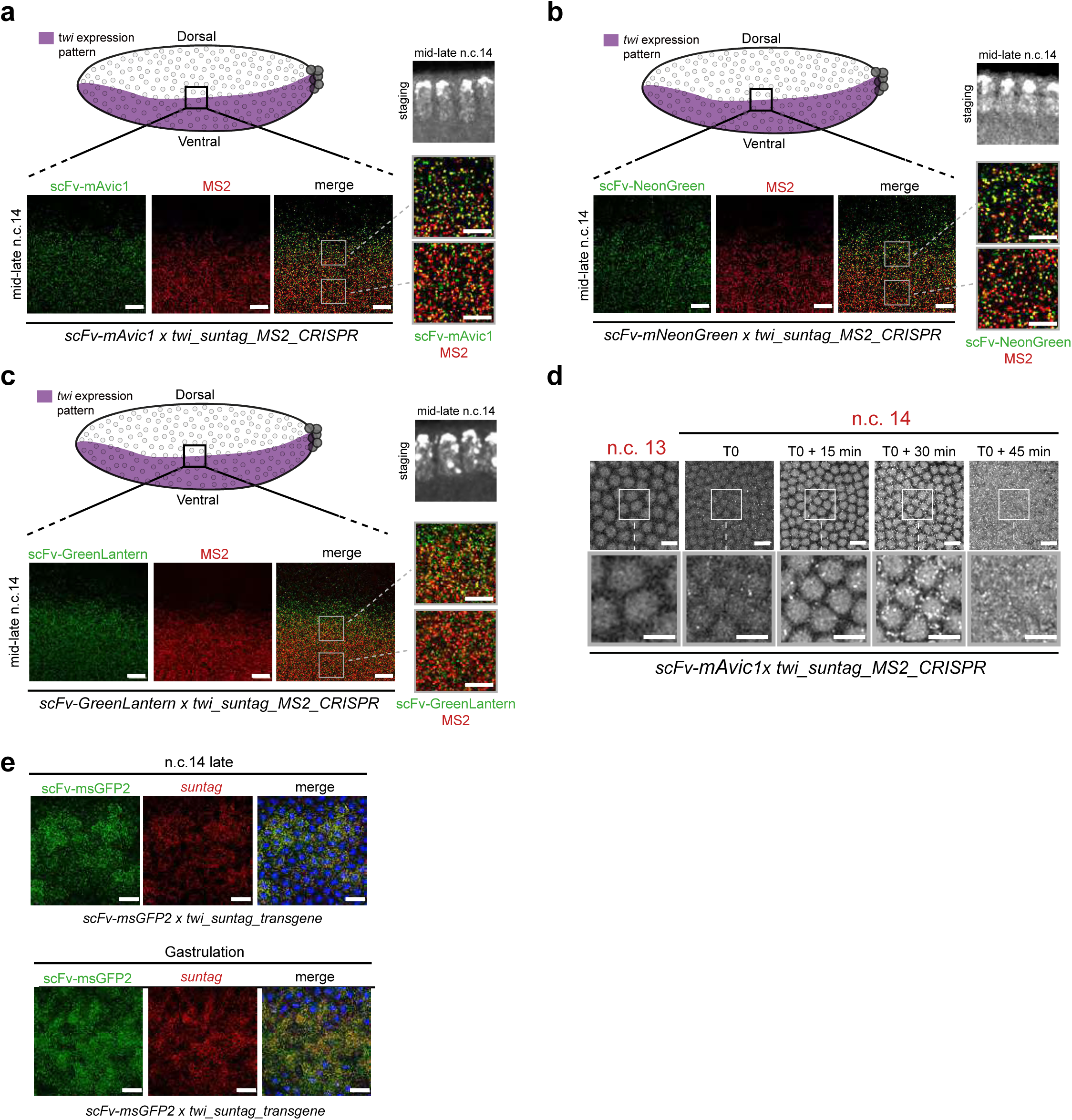
*twi* translation detection in developing embryos until gastrulation. **a.** (top) Schematic of a sagittal view of a *Drosophila* embryo with *twi* expression pattern in purple. Anterior side is on the left, posterior on the right, dorsal on the top and ventral side on the bottom. Black square represents the imaged area of the bottom panels. (bottom) A single Z-plane of confocal images from smFISH with direct fluorescent protein signal in green (scFv-mAvic1) and *MS2* probes (red) on *scFv-mAvic1 x twi_suntag_MS2_CRISPR* embryos at mid-late n.c. 14. Gray squares represent the zoomed images in the right panels. The two different zoomed images represent border (top square) and internal (bottom square) zone of the imaged pattern. Scale bars are 10µm on the larger images, and 5µm on the zoomed images. Staging is given by DAPI staining on a sagittal view of the imaged embryo (black and white image at the bottom). **b.** (top) Schematic of a sagittal view of a *Drosophila* embryo with *twi* expression pattern in purple. Anterior side is on the left, posterior on the right, dorsal on the top and ventral side on the bottom. Black square represents the imaged area of the bottom panels. (bottom) A single Z-plane of confocal images from smFISH with direct fluorescent protein signal in green (scFv-NeonGreen) and *MS2* probes (red) on *scFv-NeonGreen x twi_suntag_MS2_CRISPR* embryos at mid-late n.c. 14. Gray squares represent the zoomed images in the right panels. The two different zoomed images represent border (top square) and internal (bottom square) zone of the imaged pattern. Scale bars are 10µm on the larger images, and 5µm on the zoomed images. Staging is given by DAPI staining on a sagittal view of the imaged embryo (black and white image at the bottom). **c.** (top) Schematic of a sagittal view of a *Drosophila* embryo with *twi* expression pattern in purple. Anterior side is on the left, posterior on the right, dorsal on the top and ventral side on the bottom. Black square represents the imaged area of the bottom panels. (bottom) A single Z-plane of confocal images from smFISH with direct fluorescent protein signal in green (scFv-GreenLantern) and *MS2* probes (red) on *scFv-GreenLantern x twi_suntag_MS2_CRISPR* embryos at mid-late n.c. 14. Gray squares represent the zoomed images in the right panels. The two different zoomed images represent border (top square) and internal (bottom square) zone of the imaged pattern. Scale bars are 10µm on the larger images, and 5µm on the zoomed images. Staging is given by DAPI staining on a sagittal view of the imaged embryo (black and white image at the bottom). **d.** Snapshots each 15 minutes from movies of *scFv-msGFP2 x twi_suntag_MS2_CRISPR* embryos on the ventral side. T0 correspond to early n.c.14. White squares represent the zoomed images in the center of the panel. Note the absence of translation for *twi_suntag_MS2_CRISPR* at T0+45min, Scale bars are 10µm on the larger images, and 5µm on the zoomed images. (See related Movies S15). **e.** Single Z-planes of confocal images from smFISH with direct fluorescent protein signal in green (scFv-msGFP2) and *suntag* probes (red) on *scFv-msGFP2 x twi_suntag_transgene* embryos in late n.c. 14 (top panels) and gastrulation stage (bottom panels) on the ventral side. Nuclei are counterstained with DAPI (blue). Scale bars are 10µm.

**Figure Supp 5:**
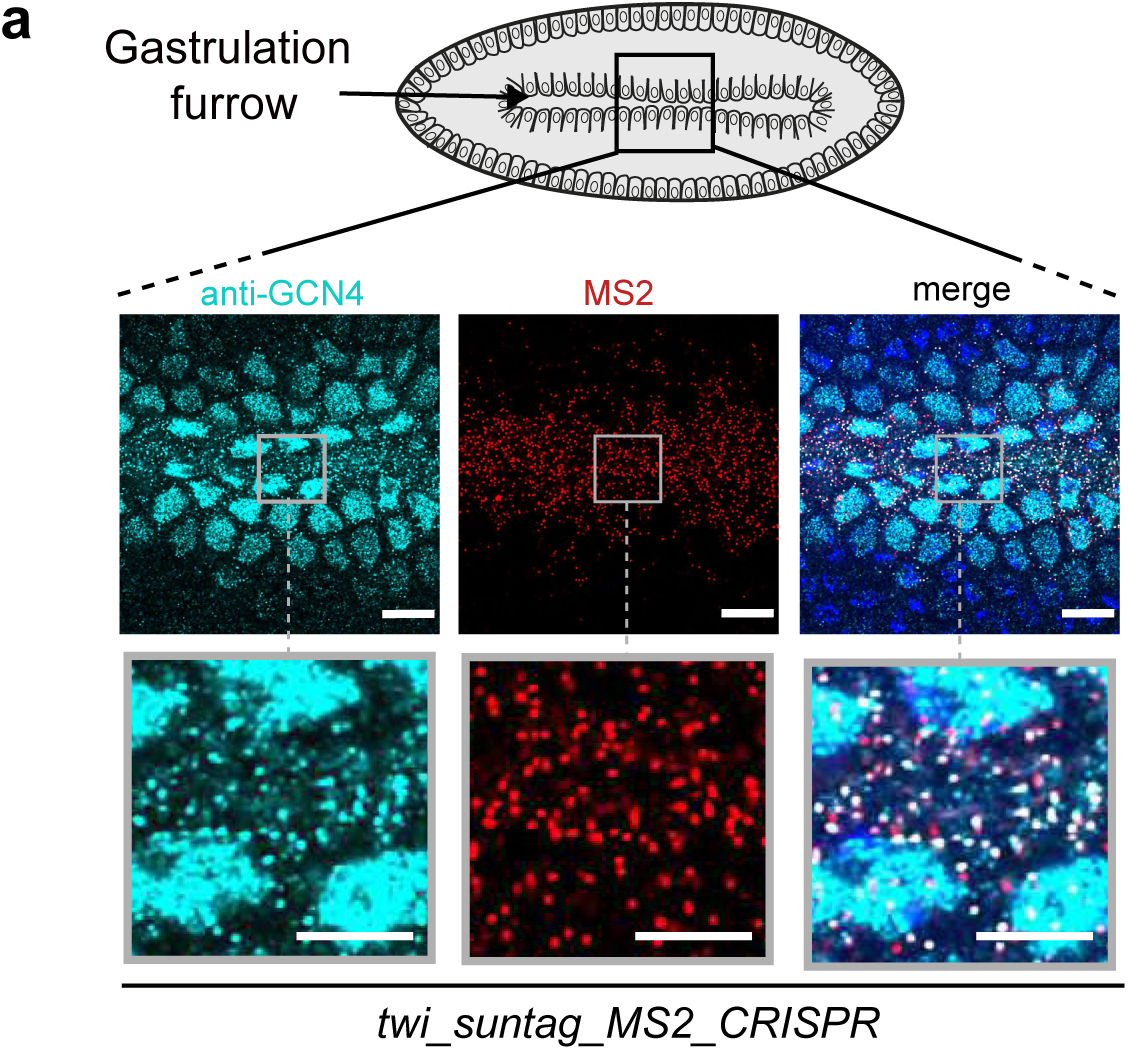
anti-GCN4 immunostaining detect translation until gastrulation. **a.** (top) Schematic of a *Drosophila* embryo on the ventral side with gastrulation furrow represented with invaginating cells. Black square represents the imaged area of the bottom panels. (bottom) Single Z-planes of confocal images from immuno-smFISH with anti-GCN4 antibody (cyan) and MS2 probes (red) on *twi_suntag_MS2_CRISPR* embryos. Grey squares represent the zoomed images in the bottom panels. Scale bars are 10µm on the larger images, and 5µm on the zoomed images.

**Figure Supp 6:**
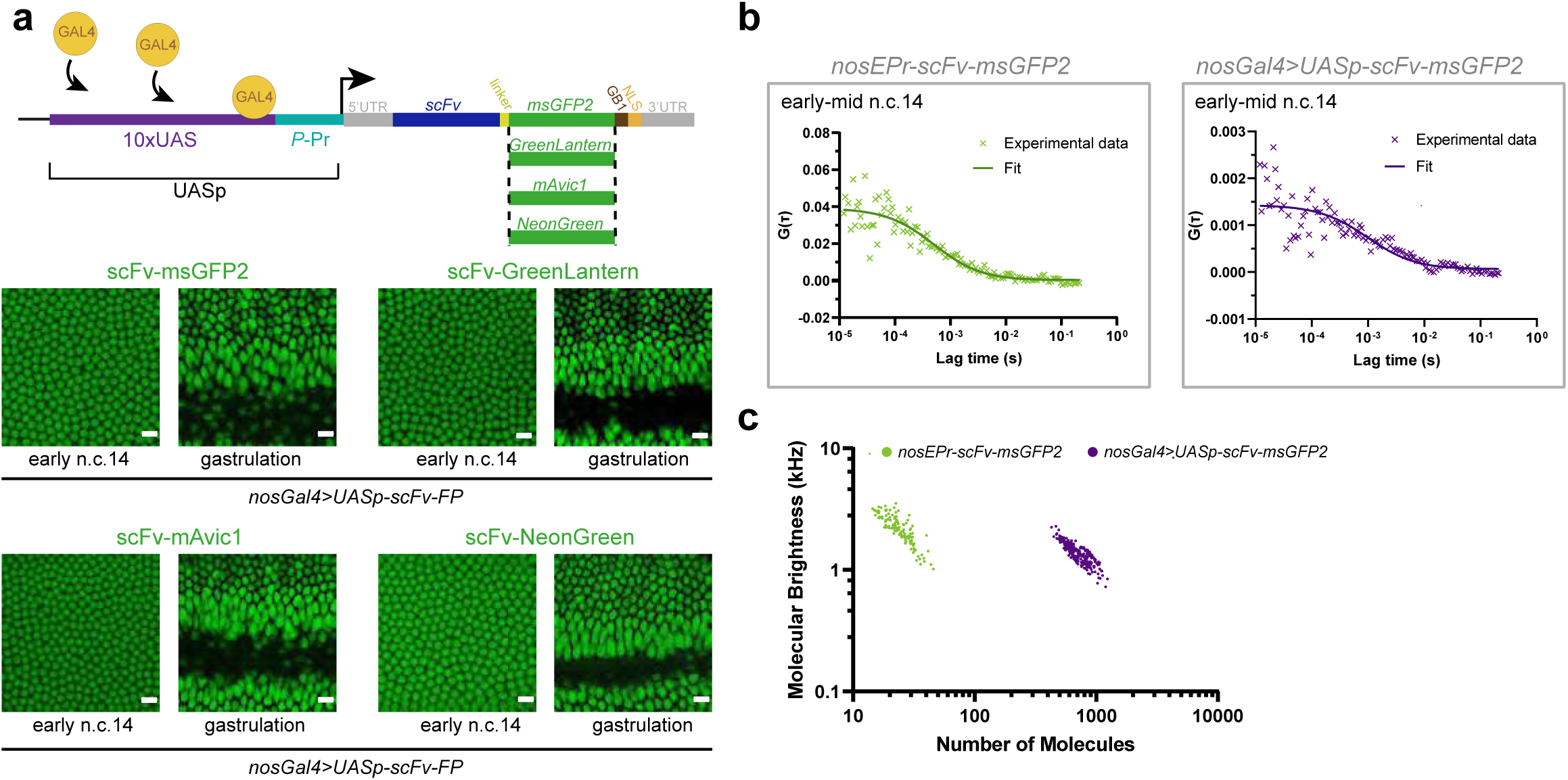
Generation of UASp-Gal4 system to overexpress scFv-Fluorescent Proteins. **a.** (top) Schematic of the different constructs expressing the different scFv-FP (scFv-GreenLantern, scFv-NeonGreen, scFv-mAvic1, or scFv-msGFP2) under the control of the UASp promoter. (bottom) Snapshots from movies of embryos carrying *UASp-scFv-msGFP2, UASp-scFv-GreenLantern, UASp-scFv-mAvic1* or *UASp-scFv-NeonGreen* and *nosGal4* at early n.c.14 and gastrulation stage. Scale bars are 10µm. (See related Movies S16-19). **b.** Representative examples of autocorrelation function from FCS performed in embryos with nosEPr-*scFv-msGFP2* at early-mid n.c. 14 stage (green) and overexpressing the scFv-msGFP2 (*nosGal4>UASp-scFv-msGFP2*) at early-mid (purple) n.c. 14 stage. Dark curves represent fitting of the data using a diffusion model (see methods). **c.** Dot plots representing the brightness as a function of the number of molecules obtained from FCS analysis (f). In embryos with nosEPr-*scFv-msGFP2* n=120, in embryos with *nosGal4>UASp-scFv-msGFP2* n=278.

**Figure Supp 7:**
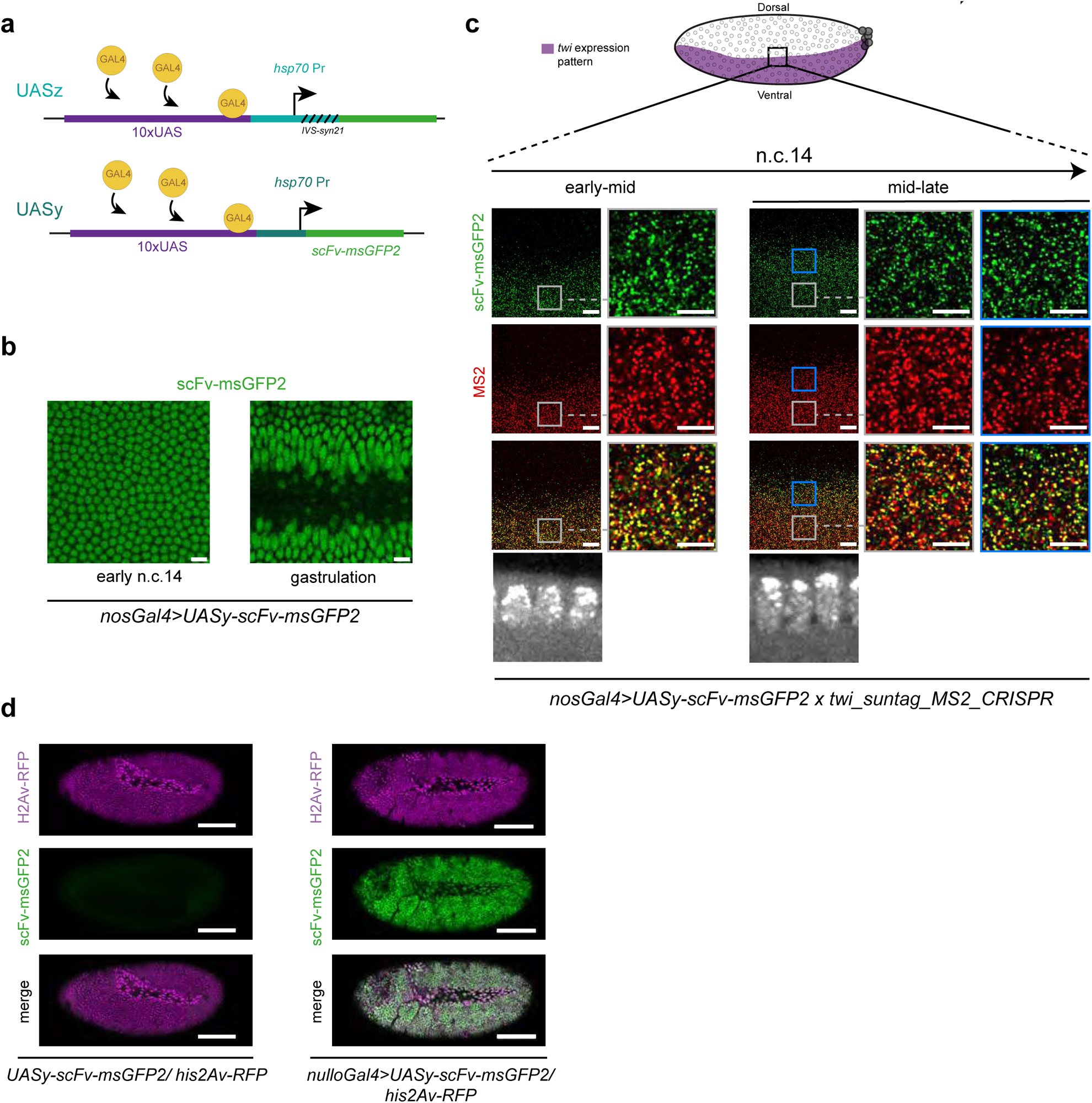
Generation of UASy-Gal4 system to express scFv-Fluorescent Proteins. **a.** Schematic of the two constructs UASz and UASy containing ten *UAS* with *hsp70* promoter and with *IVS-syn21* sequence (UASz) or without (UASy). Both of these constructs contain the *scFv-msGFP2*. **b.** Snapshots from movies of embryos carrying *UASy-scFv-msGFP2* and *nanos-Gal4* at early n.c.14 and gastrulation stage. Scale bars are 5µm. (See related Movies S21). **c.** (left) Schematic of a sagittal view of a *Drosophila* embryo with *twi* expression pattern in purple. Anterior side is on the left, posterior on the right, dorsal on the top and ventral side on the bottom. Black square represents the imaged area of the right panels. (right) A single Z-plane of confocal images from smFISH with direct fluorescent protein signal in green (scFv-msGFP2) and *MS2* probes (red) on *nosGal4>UASy-scFv-msGFP2 x twi_suntag_MS2_CRISPR* embryos in early-mid and mid-late n.c. 14. Staging is given by DAPI staining on a sagittal view of the embryo (black and white image at the bottom). Blue and white squares (border and internal zone of the pattern, respectively) represent the two different zoomed images on the right side of each panel. Scale bars are 10µm on the larger images, and 5µm on the zoomed images. **d.** Maximum intensity projected snapshots of a whole embryos confocal live tilescan expressing *UASy-scFv-msGFP2* and *Histone-H2A-RFP* (left panels) or *nulloGal4>UASy-scFv-msGFP2 and Histone-H2A-RFP* (right panels), after gastrulation. Scale bars are 100µm.

**Figure Supp 8:**
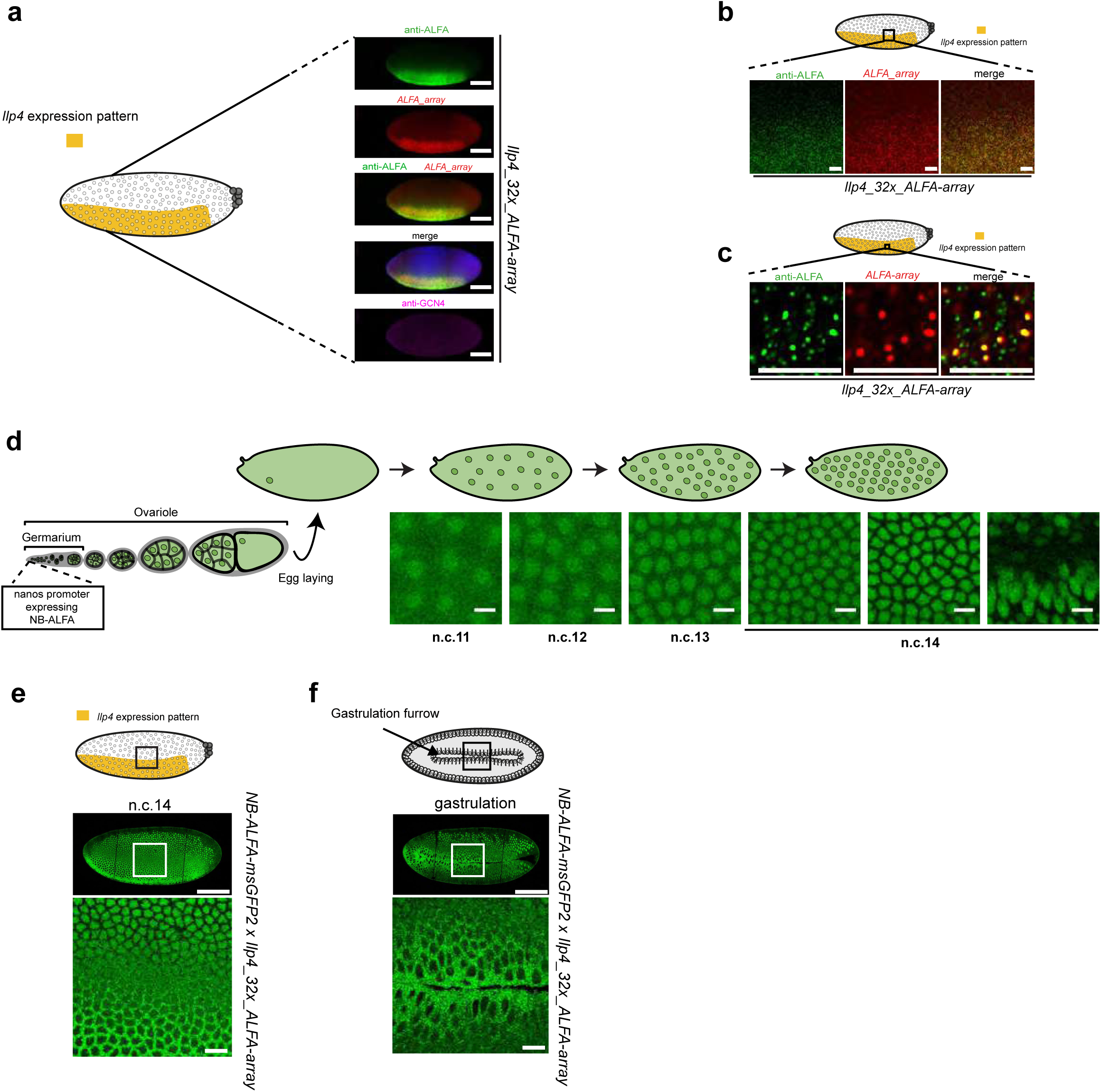
Efficient detection of *Ilp4* transgene translation with the new AlfaTag system. **a.** (left) Schematic of a sagittal view of a *Drosophila* embryo with *Ilp4* expression pattern in yellow. Anterior side is on the left, posterior on the right, dorsal on the top and ventral side on the bottom. (right) Maximum intensity projection of a whole embryo tilescan confocal images from immuno-smFISH with anti-ALFA antibody (green), *32x_ALFA-array* probes (red) and anti-GCN4 antibody (magenta) on *Ilp4_32x_ALFA-array* n.c.14 embryos. Scale bars are 100µm. **b.** (top) Schematic of a sagittal view of a *Drosophila* embryo with *Ilp4* expression pattern in yellow. Black square represents the imaged area of the bottom panels. (bottom) Single Z-planes of confocal images from immuno-smFISH with anti-ALFA antibody (green) and *32x_ALFA-array* probes (red) on *Ilp4_32x_ALFA-array* in early-mid n.c. 14 embryos at the border of *Ilp4* expression pattern. Scale bars are 10µm. **c.** (top) Schematic of a sagittal view of a *Drosophila* embryo with *Ilp4* expression pattern in yellow. Black square represents the imaged area of the bottom panels. (bottom) Single Z-planes of confocal images from immuno-smFISH with anti-ALFA antibody (green) and *32x_ALFA-array* probes (red) on *Ilp4_32x_ALFA-array* in early-mid n.c. 14 embryos to visualize single mRNA molecules in translation. Scale bars are 5µm. **d.** (top) Schematic of the expression pattern of NB-ALFA under control of the *nos EPr* during *Drosophila* oogenesis. (bottom) Snapshots from movies of embryos carrying *nos>NB-ALFA-msGFP2* in n.c. 11, 12, 13 and 14. Scale bars are 10µm. (See related Movies S23). **e.** (top) Schematic of a sagittal view of a *Drosophila* embryo with *Ilp4* expression pattern in yellow. Black square represents the imaged area of the bottom panel. (bottom) Snapshots from movies of *NB-ALFA-msGFP2 x Ilp4_32x_ALFA-array* embryos in n.c. 14. The top image is a tile scan of the whole embryo and the bottom image a zoomed area represented with a white square at the border of *Ilp4* expression pattern. Note the green signal located in nuclei outside the pattern and in the cytoplasm within *Ilp4* pattern (corresponding to tagged Ilp4 proteins). Scale bars are 100µm in the tilescan and 10µm in the zoomed image. (See related Movies S25). **f.** (top) Schematic of a sagittal view of a *Drosophila* embryo with *Ilp4* expression pattern in yellow. Black square represents the imaged area of the bottom panel. (bottom) Snapshots from movies of *NB-ALFA_msGFP2 > Ilp4_32x_ALFA-array* embryos at gastrulation stage. The top image is a tile scan of the whole embryo and the bottom image a zoomed area represented with a white square at the gastrulation furrow. Scale bars are 100µm in the tilescan and 10µm in the zoomed image. (See related Movies S24).

**Figure Supp 9:**
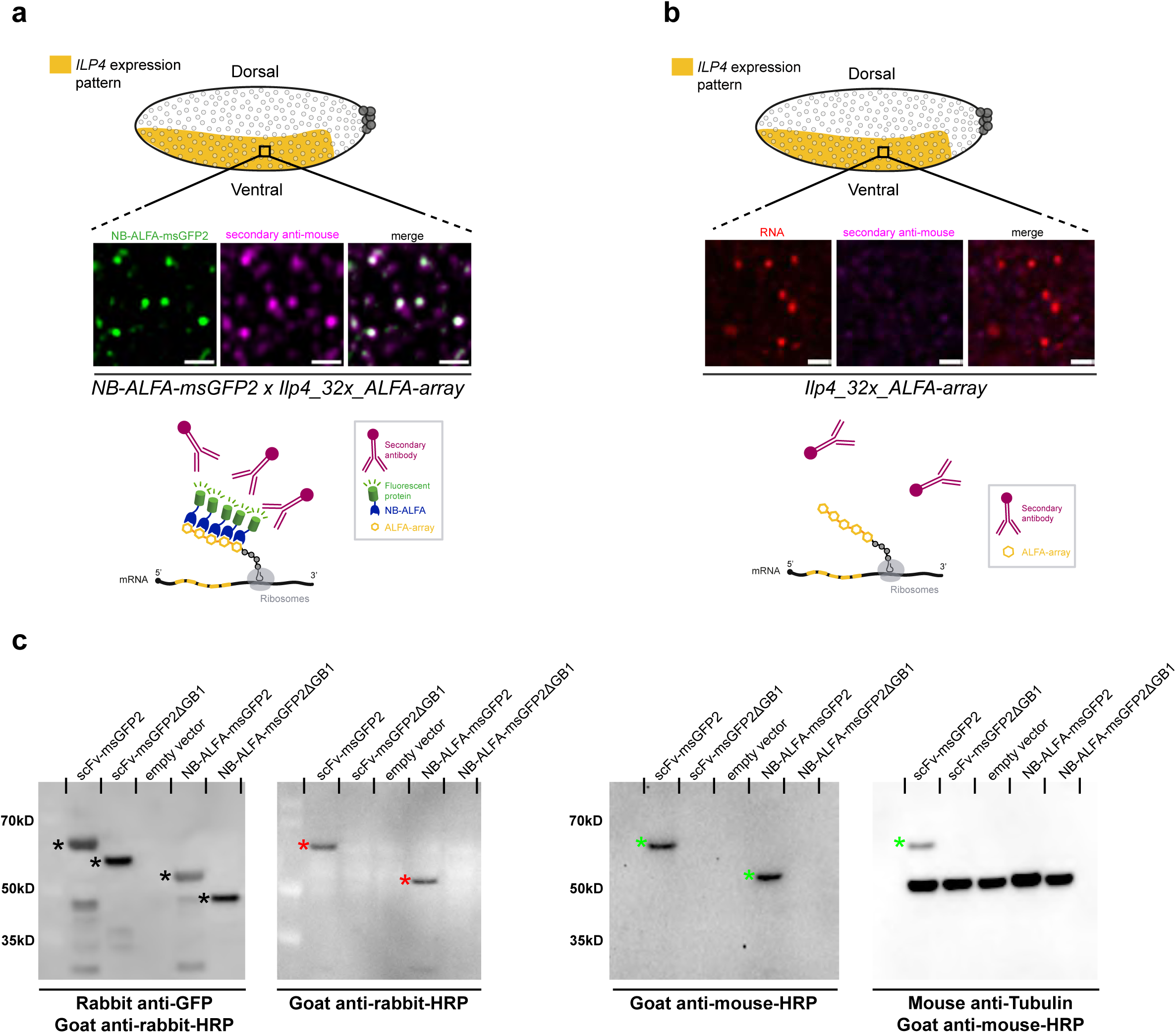
The GB1 domain is recognized by secondary antibodies. **a.** (top) Schematic of a sagittal view of a *Drosophila* embryo with *Ilp4* expression pattern in yellow. Black square represents the imaged area of the bottom panels. Single Z-planes of confocal images from immuno-smFISH with direct fluorescent protein signal in green (NB-ALFA-msGFP2) and secondary anti-mouse antibody (magenta) on *NB-ALFA-msGFP2 x Ilp4_32x_ALFA-array* embryos in n.c. 14 on the ventral side. Note the co-localization between the direct fluorescent protein signal and the secondary antibody signal as illustrated by the schematic on the bottom. Scale bars are 1µm. **b.** (top) Schematic of a sagittal view of a *Drosophila* embryo with *Ilp4* expression pattern in yellow. Black square represents the imaged area of the bottom panels. Single Z-planes of confocal images from immuno-smFISH with ALFA probes (red) and secondary anti-mouse antibody (magenta) on *Ilp4_32x_ALFA-array* embryos in n.c. 14 on the ventral side. Note that in absence of NB-ALFA the secondary antibody does not detect any signal as illustrated by the schematic on the bottom. Scale bars are 1µm. **c.** Western-blot experiments on *Drosophila* S2R+ cells transfected with scFv-msGFP2, scFv-msGFP2ΔGB1, empty vector, NB-ALFA-msGFP2 or NB-ALFA-msGFP2ΔGB1. The membrane on the left was hybridized with either anti-GFP primary and anti-rabbit secondary antibodies or with anti-rabbit secondary alone. Bands corresponding to the scFv and the NB fused to msGFP2 (with and without the GB1) are highlighted with black stars. The red stars highlight the binding of the secondary anti-rabbit-HRP to the scFv-msGFP2 and NB-msGFP2 in the presence of the GB1. The membrane on the right was hybridized with either anti-mouse secondary alone or with anti-Tubulin primary and anti-mouse secondary antibodies. The green stars highlight the binding of the secondary anti-mouse-HRP to the scFv-msGFP2 and NB-msGFP2 in the presence of the GB1.

**Figure Supp 10:**
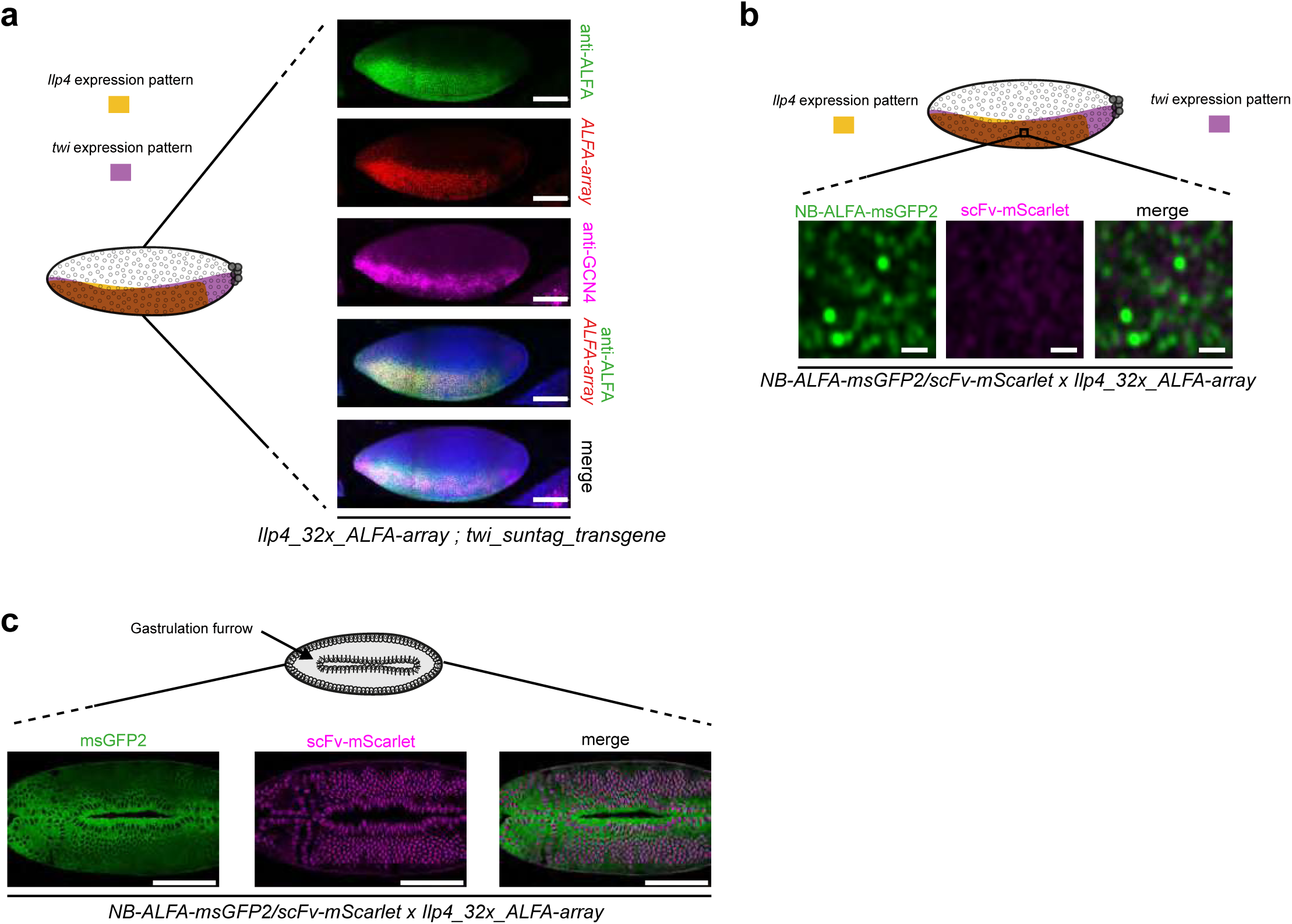
Specificity of detection of two distinct mRNA species in fixed and live samples. **a.** Maximum intensity projection of a whole embryo tilescan confocal images from immuno-smFISH with anti-ALFA antibody (green), *32x_ALFA-array* probes (red) and anti-GCN4 antibody (magenta) on *Ilp4_32x_ALFA-array; twi_suntag_transgene* embryos in n.c. 14. Scale bars are 100µm. **b.** Schematic of a *Drosophila* embryo on the ventral side with *twi* expression pattern in purple and *Ilp4* in yellow. Black square represents the imaged area of the bottom panels. One Z-plane of confocal images from live imaging of *NB-ALFA_msGFP2/scFv-mScarlet x Ilp4_32x_ALFA-array* embryos in n.c. 14. Note the absence of signal with scFv-mScarlet illustrating specificity of the scFv and the NB-ALFA. Scale bars are 1µm (see related Movies S27). **c.** (top) Schematic of a *Drosophila* embryo on the ventral side with gastrulation furrow represented with invaginating cells. Confocal snapshots of a live embryo expressing *NB-ALFA_msGFP2/scFv-mScarlet x Ilp4_32x_ALFA-array*, after gastrulation. Scale bars are 100µm.

**Figure Supp 11:**
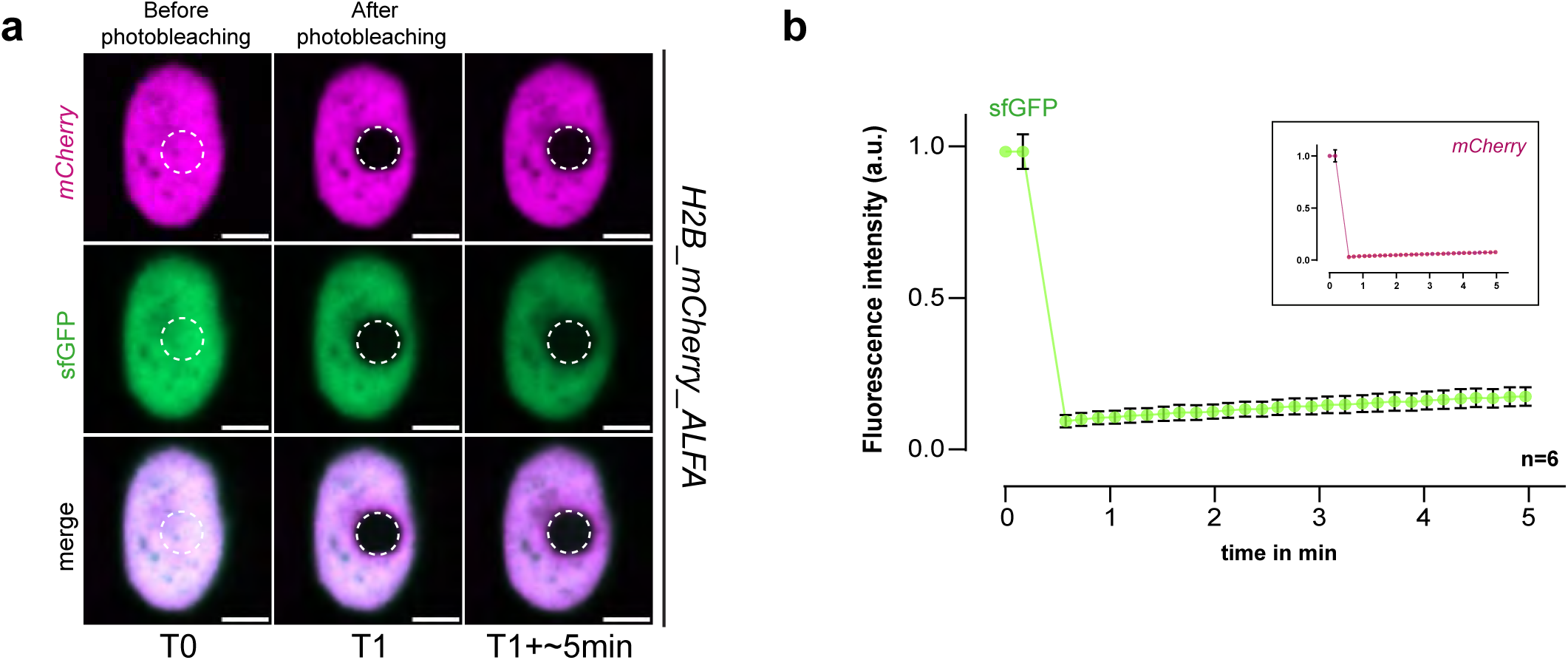
NB-ALFA exhibits a long binding time with its ALFA-Tag. **a.** Images from a FRAP experiment on HeLa cells expressing a NB-ALFA-sfGFP (green) and transfected with an H2B_mCherry_ALFAtag plasmid (magenta). Images before and after photobleaching are represented. The bleached area is represented with dashed white circles. Scale bars are 5μm. **b.** FRAP recovery curve from experiment in (c) of the fluorescence intensity as a function of time. Recovery of the NB-ALFA-sfGFP is in green and of the H2B in magenta.

## Movie Sup legends

**Movie S1**: Maximum intensity projection of confocal Z-stack live imaging of an embryo containing *scFv-sfGFP*. scale bar 10μm.

**Movie S2**: Maximum intensity projection of confocal Z-stack live imaging of an embryo containing *scFv-mAvic1*. scale bar 10μm.

**Movie S3**: Maximum intensity projection of confocal Z-stack live imaging of an embryo containing *scFv-msGFP2*. scale bar 10μm.

**Movie S4**: Maximum intensity projection of confocal Z-stack live imaging of an embryo containing *scFv-NeonGreen*. scale bar 10μm.

**Movie S5**: Maximum intensity projection of confocal Z-stack live imaging of an embryo containing *scFv-GreenLantern*. scale bar 10μm.

**Movie S6**: Maximum intensity projection of LightSheet Z-stack live imaging of *scFv-mAvic1* embryo. Upper part: dorsal view, lower part: ventral view Scale bar 100μm.

**Movie S7**: Maximum intensity projection of LightSheet Z-stack live imaging of *scFv-msGFP2* embryo. Upper part: dorsal view, lower part: ventral view Scale bar 100μm.

**Movie S8**: Maximum intensity projection of confocal Z-stack fast live imaging of an embryo containing *scFv-NeonGreen>twi_suntag_MS2_CRISPR/+.* Translation foci are in green, scale bar 1μm.

**Movie S9**: Maximum intensity projection of confocal Z-stack fast live imaging of an embryo containing *scFv-sfGFP >twi_suntag_MS2_CRISPR/+.* Translation foci are in green, scale bar 1μm.

**Movie S10**: Maximum intensity projection of confocal Z-stack fast live imaging of an embryo containing *scFv-GreenLantern >twi_suntag_MS2_CRISPR/+.* Translation foci are in green, scale bar 1μm.

**Movie S11**: Maximum intensity projection of confocal Z-stack fast live imaging of an embryo containing *scFv-msGFP2>twi_suntag_MS2_CRISPR/+.* Translation foci are in green, scale bar 1μm.

**Movie S12**: Maximum intensity projection of confocal Z-stack fast live imaging of an embryo containing *scFv-mAvic1>twi_suntag_MS2_CRISPR/+.* Translation foci are in green, scale bar 1μm.

**Movie S13**: Maximum intensity projection of confocal Z-stack live imaging of an embryo containing *scFv-msGFP2>twi_suntag_MS2_CRISPR/+.* Translation foci are in white, scale bar 10μm.

**Movie S14**: Maximum intensity projection of confocal Z-stack live imaging of an embryo containing *scFv-msGFP2>twi_suntag_transgene/+.* Translation foci are in white, scale bar 10μm.

**Movie S15**: Maximum intensity projection of confocal Z-stack live imaging of an embryo containing *scFv-mAvic1> twi_suntag_transgene/+.* Translation foci are in white, scale bar 10μm.

**Movie S16**: Maximum intensity projection of confocal Z-stack live imaging of an embryo containing *nos-Gal4>UASp-scFv-GreenLantern/+*. scale bar 10μm.

**Movie S17**: Maximum intensity projection of confocal Z-stack live imaging of an embryo containing *nos-Gal4>UASp-scFv-msGFP2/+*. scale bar 10μm.

**Movie S18**: Maximum intensity projection of confocal Z-stack live imaging of an embryo containing *nos-Gal4>UASp-scFv-NeonGreen/+*. scale bar 10μm.

**Movie S19**: Maximum intensity projection of confocal Z-stack live imaging of an embryo containing *nos-Gal4>UASp-scFv-mAvic1/+*. scale bar 10μm.

**Movie S20**: Comparison of maximum intensity projection of confocal Z-stack live imaging of an embryo containing *scFv-msGFP2>twi_suntag_MS2_CRISPR/+ or nos-Gal4>UASp-scFv-mAvic1; twi_suntag_MS2_CRISPR/+*. Translation foci are in white. Translationnal arrest is seen *scFv-msGFP2>twi_suntag_MS2_CRISPR/+ but not* in *nos-Gal4>UASp-scFv-mAvic1; twi_suntag_MS2_CRISPR/+*. scale bar 10μm.

**Movie S21**: Maximum intensity projection of confocal Z-stack live imaging of an embryo containing *nos-Gal4>UASy-scFv-msGFP2/+*. scale bar 10μm.

**Movie S22**: Maximum intensity projection of confocal Z-stack live imaging of an embryo containing *nullo-Gal4>UASy-scFv-msGFP2/+*. scale bar 100μm.

**Movie S23**: Maximum intensity projection of confocal Z-stack live imaging of an embryo containing *NB-ALFA-msGFP2*. scale bar 10μm.

**Movie S24**: Maximum intensity projection of confocal Z-stack live imaging of an embryo containing *NB-ALFA-msGFP2> Ilp4_32x_ALFA-array/+*. Scale bar 10μm.

**Movie S25**: Maximum intensity projection of confocal tile-scan with Z-stack live imaging of an embryo containing *NB-ALFA-msGFP2> Ilp4_32x_ALFA-array/+*. Scale bar 100μm.

**Movie S26**: Single plane of a confocal live imaging of an embryo containing *twi_suntag_MS2_CRISPR/+; NB-ALFA_msGFP2/scFv-mScarlet x Ilp4_32x_ALFA-array* during n.c.14. Scale bar 5μm.

**Movie S27**: Single plane of a confocal live imaging of an embryo containing *NB-ALFA_msGFP2/scFv-mScarlet x Ilp4_32x_ALFA-array* during n.c.14. Scale bar 5μm.

## Notes

### Competing Interest Statement

The authors have declared no competing interest.

### Summary of Updates

We have revised the manuscript to improve readability and rearranged the figures for better clarity. These modifications aim to enhance the overall understanding and flow of the manuscript.

